# State-dependent aperiodic EEG dynamics track cortical network reorganization in chronic epilepsy

**DOI:** 10.64898/2026.08.20.745947

**Authors:** Garima Chauhan, Kshitij Kumar, Deepti Chugh, Subramaniam Ganesh, Arjun Ramakrishnan

## Abstract

Aperiodic (1/f-like) EEG activity has rapidly become a popular noninvasive marker of cortical network state, proposed to index excitation–inhibition (E/I) balance and increasingly applied across neurological and psychiatric disorders. However, whether this approach remains reliable in the pathological brain, where disease progressively reorganizes neural networks, alters signal morphology, and drives continuous transitions between cortical states has yet to be systematically established. Using a medication-free genetic model of chronic epilepsy (Lafora disease; Epm2a−/− mice), we tracked the aperiodic component of the cortical EEG across resting wakefulness, isoflurane anesthesia, and PTZ-induced seizures of graded severity, asking how a single spectral marker behaves as the brain moves between states. Epileptic mice exhibited systematically steeper aperiodic exponents than controls, an effect that persisted after removal of interictal epileptiform discharges and was replicated using independent time-resolved spectral parameterization. Slopes steepened predictably under GABAergic anesthesia, supporting the interpretation that aperiodic activity captures biologically meaningful state transitions beyond simple contamination by pathological waveforms. Across seizure phases, the aperiodic exponent varied systematically, however, the exponent flattened during ictal activity in step with the dominant discharge morphology, revealing that pathological waveform shape itself is a substantial contributor to seizure-state exponent changes. Together, these findings indicate that aperiodic EEG dynamics reflect a combination of chronic network-state reorganization and waveform-shape-driven spectral distortion, with their relative contributions varying across brain states. These results support spectral parameterization as a sensitive approach for tracking pathological neural activity in chronic epilepsy while delineating its interpretive boundaries in the presence of pathological waveforms.

**Significance statement:** The “aperiodic” structure of brain electrical activity has rapidly become a popular noninvasive marker of cortical state, widely used to infer the balance of excitation and inhibition in health and disease. But how faithfully does it report on a brain that is both diseased and continually changing state? Using a genetic mouse model of chronic epilepsy, we show that this marker conflates two distinct phenomena: lasting reorganization of cortical networks, and distortion caused by the shape of pathological discharges themselves. Separating these contributions across seizure stages and severities yields a more accurate and cautious framework for interpreting this EEG signature; one that matters broadly, as spectral analysis is increasingly applied to noisy, disease- and treatment-altered recordings in neurological and psychiatric patients.

## 1.1 Introduction

Epilepsy offers a powerful model for studying how the pathological brain reorganizes progressively over time, as recurrent seizures drive measurable, evolving changes in brain-wide connectivity and cortical excitability (Liu et al., 2023; Barrios-Martinez et al., 2025). Epileptogenesis involves ongoing homeostatic adaptations in excitation–inhibition (E/I) balance that evolve over time and across seizure states (Žiburkus et al., 2013; Chen et al., 2022; Xie et al., 2024). Repeated seizures trigger long-lasting neuroplastic changes (Wang et al., 2025), destabilize local and global functional organization, and increase vulnerability to subsequent events (Liang et al., 2020). Consequently, epilepsy is now understood as a progressive disorder in which alterations in network architecture and neuronal excitability continuously reshape brain function.Understanding these evolving network dynamics is central to explaining seizure initiation, propagation, and recovery. However, despite growing evidence that epileptogenesis is accompanied by continuous reorganization of neural circuits, how large-scale cortical network organization evolves throughout chronic epilepsy and across different seizure stages and severities remains poorly understood. This gap is particularly pronounced in human studies, where ethical constraints and the widespread use of anti-epileptic drugs (AEDs) obscure intrinsic circuit dynamics (Belete, 2023).

The inhibitory dynamics of epilepsy make it an unusually stringent testbed for any marker of cortical excitability, precisely because inhibition operates paradoxically,varying across brain and disease stages rather than moving in a single direction.. This paradoxical, multifaceted role of inhibition is well documented:absence seizures exhibit stronger inhibitory tone and reduced E/I ratios (Eisenstein et al., 2024).While impaired inhibition contributes to hyperexcitability, enhanced inhibition can also precede seizure onset, shape synchronization and contribute to seizure termination. (Shao et al., 2019; Xie et al., 2024; Žiburkus et al., 2013). Consequently, E/I balance is increasingly viewed as a dynamic network property rather than a fixed characteristic of epileptic cortex (Muthukumaraswamy and Liley, 2018), yet how these processes manifest at the systems level in vivo remains unclear.

Against this backdrop of dynamic, state-dependent inhibition,aperiodic EEG activity, once regarded as nonspecific neural noise, is now increasingly recognized as a physiologically meaningful signal and has emerged as a candidate biomarker across multiple neurological and psychiatric disorders (Phillips et al., 2024; Woronko et al., 2025, Aggarwal & Ray., 2025). The aperiodic (1/f) component of the neural power spectrum has been proposed as a systems-level descriptor related to cortical excitability, with computational models with in vivo validation in rodents, linking the spectral slope to shifts in excitation and inhibition (Gao et al., 2017; Chini et al., 2022).It is best understood, however, as consistent with, rather than a direct measurement of, underlying synaptic E/I ratios. Because the exponent is inexpensive to compute from virtually any recording and is proposed to index such a fundamental circuit property, it has been adopted rapidly across sleep, anesthesia, development, and a widening range of neurological and psychiatric conditions (Lendner et al., 2020; Höhn et al., 2024; Ostlund et al., 2022; Woronko et al., 2025).. This rapid uptake, however, has outpaced systematic tests of how faithfully the exponent reports on brains that are simultaneously diseased and shifting between cortical states, precisely the conditions under which it is most often applied.

Within epilepsy specifically, the aperiodic exponent has been shown to vary dynamically with disease state, rising prior to seizures, remaining elevated during ictal activity, and decreasing following responsive neurostimulation, pointing to its potential utility as a state-sensitive neural biomarker in clinical settings. Several studies report that the aperiodic slope steepens under inhibitory conditions such as sleep or anesthesia (Lendner et al., 2020; Höhn et al., 2024) and flattens during heightened excitation (Gao et al., 2017). A complementary spectral offset parameter provides insight into broadband excitability (Donoghue et al., 2020). Steepening of the aperiodic slope has been reported during ictal states (Liu et al., 2023) and in temporal lobe epilepsy (Duma et al., 2024), though its relationship to seizure phase, severity, and underlying network reorganization remains unresolved.

A critical and underappreciated challenge, however, concerns the interpretive specificity of the aperiodic exponent in pathological recordings.Recent work has directly demonstrated that the shape of epileptic discharges substantially influences the estimated aperiodic exponent, such that observed slope changes during seizure states may reflect waveform morphology rather than, or in addition to, underlying shifts in network excitability (Heidiri et al., 2025). Consequently, an important unresolved question is whether changes in the aperiodic exponent primarily reflect chronic disease-related network reorganization, transient state-dependent changes in inhibition and excitation, pharmacological modulation of cortical dynamics, or pathological waveform morphology. Resolving these possibilities is essential if spectral parameterization is to provide mechanistic insight or serve as a reliable biomarker of epilepsy.

Compounding this challenge, uncertainty in interpreting the aperiodic exponent extends beyond pathological waveform morphology, because periodic and aperiodic spectral components are themselves not independent. harmaco-EEG and MEG studies suggest that changes in canonical frequency bands may partly reflect alterations in the aperiodic component (Donoghue et al., 2020; Ostlund et al., 2022), which is itself sensitive to GABAergic modulation (Muthukumaraswamy et al., 2013; Stock et al., 2020). GABA-A receptor subtype plasticity (Fritschy, 2008; Mathew et al., 2012) and inhibitory network dynamics underlying beta oscillations (Yamawaki et al., 2008; Rossiter et al., 2014) further highlight the need for analytical approaches that jointly characterize periodic and aperiodic features when probing cortical network dynamics.

We therefore asked not whether the aperiodic exponent provides a direct measure of excitation–inhibition balance, but under what physiological conditions it provides meaningful information about cortical network organization. To address this question, we combined a chronic genetic model of epilepsy with pharmacological manipulation and spectral parameterization to characterize state-dependent cortical network dynamics across multiple biologically distinct conditions. Because Lafora disease is characterized by progressive epileptogenesis in the absence of chronic anti-epileptic treatment, it provides an opportunity to examine intrinsic network reorganization across disease progression. In our study, we used the Epm2a−/− (laforin knockout) mouse model of Lafora disease, a medication-free, genetically defined, and progressively symptomatic model.

Using bilateral screw EEG recordings from wild-type and Epm2a−/− mice, we quantified aperiodic and oscillatory spectral features during resting wakefulness, GABA-A receptor modulation with isoflurane anesthesia, and PTZ-induced seizures spanning preictal, ictal, postictal, and different seizure severity states. This design allowed us to ask two closely related questions. First, whether the aperiodic exponent provides a robust marker of network state that remains stable across interictal discharges, alternative parameterization methods, and pharmacological perturbation. Second, where its interpretation becomes constrained because pathological waveform shape emerges as a major contributor to spectral slope estimates.

Establishing both the robustness and the boundary conditions of this marker in a controlled model carries direct implications for the growing number of studies applying spectral parameterization to noisy, disease- and treatment-altered recordings across EEG, MEG, and intracranial electrophysiology. More broadly, this framework provides a systems-level test of when aperiodic activity tracks evolving cortical network states, and when its interpretation is constrained by pathological waveform distortions.

## 2. Methods/Methodology

Detailed procedures, algorithm parameters, and full statistical model specifications for the analyses below are provided in Supplementary Methods.

### 2.1 Experimental Animals

Laforin-deficient mice represent a validated animal model of Lafora disease, a fatal progressive myoclonus epilepsy characterised by abnormal glycogen accumulation and neurodegeneration (Parihar and Ganesh, 2024). The Epm2a−/− mouse model of Lafora disease was generated as described previously (Ganesh et al., 2002). Recordings were performed in mice aged 6–8 months, a timepoint at which the Epm2a−/− line exhibits well-established Lafora disease pathology, including polyglucosan (Lafora) body accumulation, neurodegenerative changes, and spontaneous epileptic activity, consistent with the progressive phenotype characterized across age in this model (Burgos et al., 2023). The terms *laforin knockout (LKO)* and *Epm2a−/−* are used interchangeably throughout the manuscript. All experiments were performed in accordance with the institutional animal ethics committee (IAEC) guidelines after obtaining approvals for the protocols (IAEC project number: IITK/IAEC/2021/1137). Mice were bred in the CEAF (Central Experimental Animal Facility), at IIT Kanpur. Both male and female mice were used for the study. Animals were group-housed in standard cages under conventional laboratory conditions and individually housed from the day of electrode implantation onward. Environmental conditions were maintained at 22 ± 2 °C temperature, 55 ± 5% humidity, and a 12 h light/dark cycle (lights on at 7 a.m.). Food and water were provided *ad libitum*.

### 2.2 Surgical Protocol, Animal Preparation

All surgical procedures were conducted on both wildtype and transgenic C57BL/6 mice (Laforin knockout model) using aseptic techniques. Four active screw EEG electrodes were implanted stereotactically over the frontal and temporal cortices, with ground and reference electrodes placed over the cerebellum. Coordinates relative to Bregma were: right frontal (AP +2.5 mm, ML +1.3 mm), left frontal (AP +2.5 mm, ML −1.3 mm), right temporal (AP −3.0 mm, ML +2.5 mm), left temporal (AP −3.0 mm, ML −2.5 mm), and reference (AP −6.0 mm, ML 0.0 mm).

### 2.3 EEG Acquisition

Electrophysiological recordings were carried out in freely moving mice using the Intan RHS stim/recording system (Intan Technologies, LLC). Animals were connected via a lightweight and flexible tether to minimize movement restriction. To probe how anesthetic modulation of cortical state affects the aperiodic (1/f) component of the EEG, we compared baseline recordings with recordings obtained under 1.5% isoflurane in wild-type (WT) and Epm2a−/− (LKO) mice. Isoflurane was chosen because GABA-A receptor–acting anesthetics typically steepen the spectral slope (Lendner et al., 2020; Muthukumaraswamy & Liley, 2018). EEG traces were visually monitored for burst-suppression activity throughout each recording to confirm comparable anesthesia depth across animals and genotypes.

### 2.4 PTZ dose

PTZ is an epileptogenic compound used to induce epilepsy and measure neuronal excitability.For the induction of seizures, a single dose of PTZ 55 mg/kg (convulsive) (Sigma-Aldrich™, St. Louis, U.S.A.; purity ≥ 99%) was administered intraperitoneally.

### 2.5 Seizure grading and waveform classification

Seizure severity in PTZ-induced mice was assessed using a modified Racine scale synchronized with EEG/video recordings (Van Erum et al., 2019). Behavioral seizures were classified as low-grade (scores 0–2; partial/focal seizures), high-grade (scores 3–4; generalized seizures with bilateral motor involvement and loss of postural stability), and severe tonic–clonic seizures (scores 5–7; sustained clonus, tonic posturing, and possible loss of righting reflex). Behavioral scoring was performed by trained observers blinded to experimental groups. For tonic–clonic seizures, 30 s epochs were extracted from the preictal period (immediately before electrographic seizure onset), ictal period (sustained electrographic seizure with corresponding behavioral manifestations), and postictal period (immediately after seizure termination, representing EEG depression). Grade-2 and grade-4 seizure epochs were selected only when behavioral scores matched corresponding electrographic abnormalities. Epochs containing state transitions, insufficient duration, or excessive artifacts were excluded.

Pathological EEG waveforms were classified into four electrographic patterns: spike-wave discharges (SWD), polyspike-burst discharges (PSB), high-amplitude polyspikes (HAPS), and low-amplitude high-frequency rhythmic waveforms (LAHFRW), based on established morphological criteria. SWDs were associated with non-motor seizures, whereas HAPS and LAHFRW were characteristic of severe tonic–clonic seizures, with LAHFRW typically followed by postictal EEG depression. High-amplitude polyspike and spike-wave patterns accompanied the most severe behavioral convulsions, while prolonged high-amplitude polyspiking with subsequent high-frequency oscillations specifically distinguished the most severe tonic–clonic seizures from lower-amplitude discharge patterns associated with milder seizure stages (Van Erum et al., 2019; Lüttjohann et al., 2009).

### 2.6 EEG Data Processing and Analysis

All offline data processing and analysis were initiated in Brainstorm software (Tadel et al., 2011), running in MATLAB (version R2024a; The MathWorks Inc., Natick, MA, USA).

#### 2.6.1 Fitting Oscillations & One-Over-F’ (FOOOF) algorithm

To decompose the power spectral density (PSD) of clean epochs into periodic and aperiodic components, we used the ’Fitting Oscillations & One-Over-F’ (FOOOF) algorithm (Donoghue et al., 2020), implemented in Python. This separation is crucial as traditional band-power measures can conflate these two signals (Donoghue et al., 2020). The FOOOF algorithm models the aperiodic component as a 1/f function and identifies periodic components as Gaussian peaks rising above the aperiodic slope.

#### 2.6.2 Interictal epileptiform discharges (IEDs) detection

To detect IEDs we use a literature-matched approach (Kopf et al., 2024), signals were bandpass filtered between 25 and 80 Hz using a fourth-order Butterworth filter. The analytic amplitude of the filtered signal was computed via the Hilbert transform and z-score normalized relative to the artifact-free segments of the recording. IED events were identified where this normalized analytic amplitude exceeded a threshold of three standard deviations (z > 3) for a continuous duration of 20 to 100 ms. Event timestamps were time-locked to the local peak of the analytic amplitude, and any candidate event coinciding with a flagged artifact segment was discarded.

#### 2.6.3 Time-Resolved Spectral Parameterization (SPRiNT)

Spectral parameterization of the EEG signal was performed using SPRiNT (Spectral Parameterization Resolved in Time; Wilson et al., 2022)., implemented within Brainstorm as process_sprint. SPRiNT extends the FOOOF/specparam framework (Donoghue et al., 2020) to resolve aperiodic and periodic spectral features across time by applying the specparam algorithm to a series of short-time Fourier transform (STFT) windows within each epoch.

For each 30-second epoch, a sliding STFT was computed using windows of 1 second duration with 90% overlap, producing a time-frequency representation with a frequency resolution of 1 Hz.

#### 2.6.4 Data Quality Control

Prior to statistical analysis, data quality was ensured through two filtering steps. First, only epochs with FOOOF model fits achieving an R² > 0.8 were retained, ensuring high-quality spectral parameterization. Second, to remove outliers in the aperiodic exponent, we excluded epochs where the exponent deviated by more than 2 standard deviations from the animal-specific mean, calculated separately for each animal.

### 2.7 Statistical Analysis

All statistical analyses were conducted using Bayesian multilevel regression models implemented in the brms package (Bürkner, 2017) in R (version 4.2.1; R Core Team, 2024). Bayesian approaches were chosen for their ability to account for hierarchical data structures, provide full posterior distributions for parameter estimates, and quantify uncertainty in a probabilistic framework (Kruschke, 2015). We used separate comparison group-wise since not all animals had grade-4 or TC after the same dose of PTZ. So changes during different grades were compared to baselines of the same subject to rule out group level effects.

#### 2.7.1 Effect Size Calculation

For all models, effect sizes were calculated as standardized coefficients by dividing each Stage coefficient (β_Stage) by the residual standard deviation (σ) from the model:Cohen’s d = β_Stage / σ. This approach yields standardized effect sizes representing the magnitude of stage differences relative to within-unit residual variation, independent of measurement scale. These standardized coefficients were summarized using posterior means and 95% credible intervals.

#### 2.7.2 Data availability

EEG datasets supporting the findings of this study have been deposited in Figshare and are available at https://doi.org/10.6084/m9.figshare.33038867.

#### 2.7.3 Code availability

We deposited the code of this project in the project repository and made it openly available and licensed for reuse at github.com/kshtjkumar/E_I_activity_epilepsy.git.

## 3. Results

### 3.1 Steeper resting aperiodic exponents in epileptic Epm2a−/− mice persist independently of interictal epileptic discharges

Understanding how changes in aperiodic spectral dynamics reflect seizure vulnerability requires separating aperiodic and oscillatory contributions to cortical activity. In Epm2a−/− mice, previous work has documented a reduction in cortical GABAergic neurons and altered inhibitory tone (Ortolano et al., 2014). To assess how these cellular changes manifest at the systems level, we analyzed resting-state wake EEG using spectral parameterization to dissociate aperiodic and periodic components and compare them across Laforin knockout (LKO) and wild-type (WT) animals. Based on the reported loss of GABAergic neurons in neonatal LKO mice, we initially expected flatter aperiodic slopes and elevated beta activity.

We first examined aperiodic parameters during wakeful rest baseline recording from both genotypes. Subject-specific aperiodic fits were derived from individually estimated exponent and offset values, with representative WT and LKO traces illustrating higher epileptiform discharges in knockout mice (Fig. 1A–B; stats provided, Fig. S1.). Fig. 1C is the representative power spectrum parameterized with FOOOF, illustrating the aperiodic component (broadband offset and exponent; dashed line) and the periodic component (central frequency and relative power above the aperiodic fit). Grey shading marks the canonical beta band (12–30 Hz). Contrary to our initial hypothesis, Epm2a−/− animals showed significantly steeper aperiodic exponents (1.35 ± 0.31) relative to WT mice (1.10 ± 0.23) (Fig. 1D; n = 12, and 48 sessions (30-min each); β = 0.30, 95% CI [0.09, 0.52]; effect size = 1.58, 95% CI [0.48, 2.70]). No interaction effect was observed.

**Figure 1.**
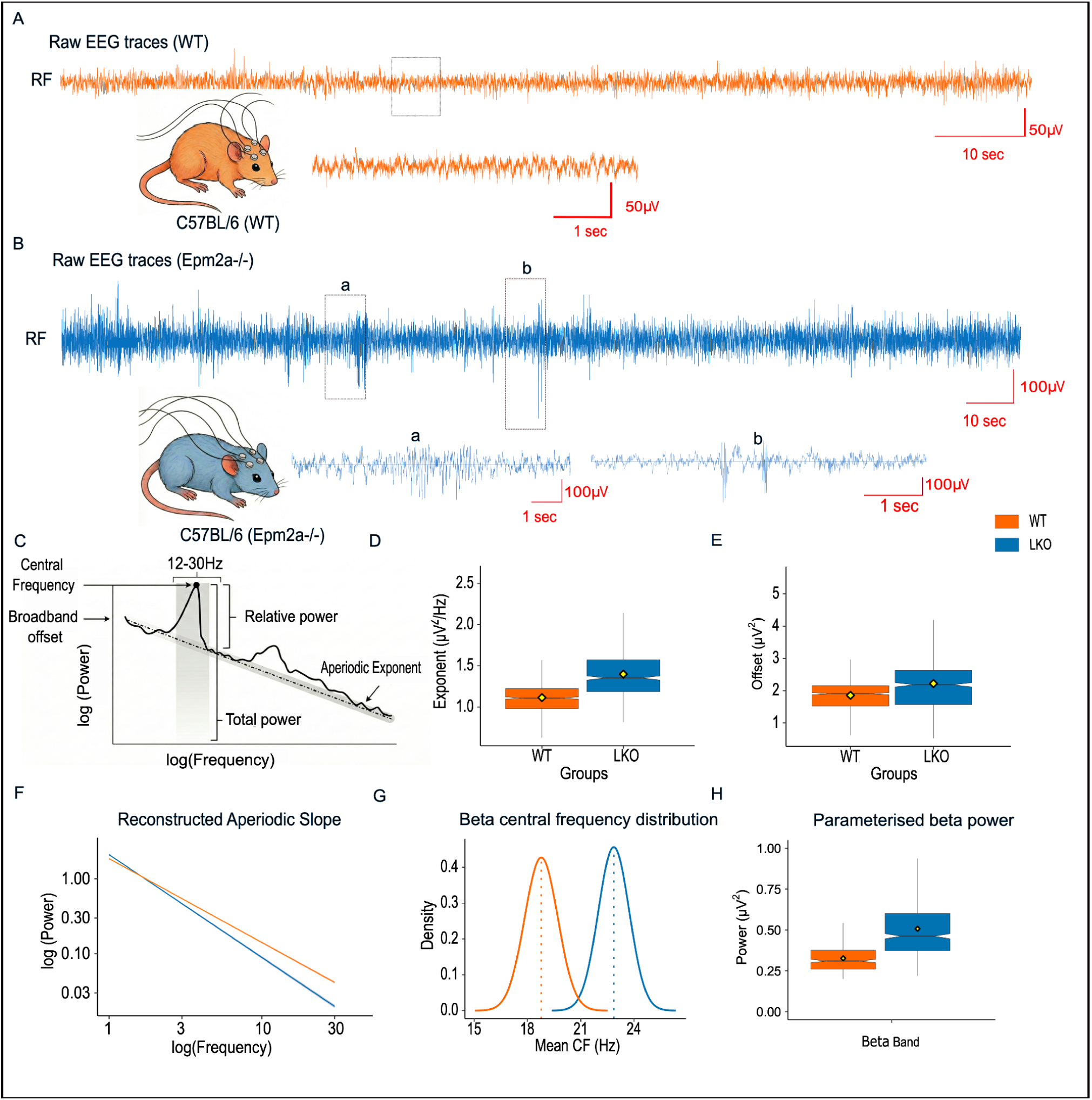
Steeper resting aperiodic exponents distinguish epileptic from wild-type mice. **A.** Intracranial neurophysiological activity was recorded using screw EEG in C57BL/6 mice at age of 6-8 months. Shown are representative artifact-free baseline awake traces (orange), resampled at 2000 Hz, displayed at a scale of 50 μV/10 sec, and zoomed at 50 μV/1 sec. **B.** Representative raw EEG traces from knockout Epm2a−/− mice (blue), resampled at 2000 Hz, showing increased epileptic discharges, displayed at 100 μV/10 sec and zoomed at 100 μV/1 sec, respectively. **C.** Representative neural power spectrum parameterized using the FOOOF algorithm, illustrating periodic components (central frequency, relative power) and aperiodic components (exponent, offset), with a prominent beta-band peak in the canonical 12–30 Hz range (grey shaded region).The offset (y-intercept) and slope (exponent) define the aperiodic component in log-transformed power. **D.** Mean aperiodic exponent (slope) compared between both WT (C57BL/6) and KO (Epm2a−/− , LKO)mice. **E.** Mean aperiodic offset (y-intercept) compared between both WT (C57BL/6) and KO (Epm2a−/− , LKO)mice. **F.** Reconstructed aperiodic slope showed steepening in LKO animals at 30s epoch from 1 to 30hz frequency range when compared between genotypes. **G.** Periodic Central Frequency (Hz) peak in beta frequency band was higher in Epm2a−/− mice compared to WT mice. **H.** Aperiodic adjusted power in beta frequency (12–30 Hz) was significantly higher in Epm2a−/− mice compared to WT mice. Box plot represents median/IQR, notches are the 95% CI of the median, and the diamond is the mean. WT (C57BL/6) represented with orange and KO (Epm2a−/− , LKO)mice represented with blue colour in this figure.

Critically, this steepening persisted after excluding IEDs: SPRiNT decomposition showed that IED-containing windows made up only 4.8% of the LKO baseline, and the exponent recomputed from the remaining IED-free windows (95.2%) remained steeper in LKO than WT (**Fig. S1**) In contrast to the exponent, the aperiodic offset differed significantly between genotypes, with LKO animals showing a higher offset than WT (2.24 ± 0.95 vs. WT: 1.86 ± 0.47; Fig. 1E; β = 0.81, 95% CI [0.05, 1.54]; effect size = 1.77, 95% CI [0.12, 3.36]), indicating greater broadband spectral power in the epileptic cortex. This genotype difference further interacted with recording region (LKO × temporal β = −0.66, 95% CI [−0.75, −0.56]; effect size = −1.43, 95% CI [−1.65, −1.22]), indicating that the offset elevation in LKO animals was regionally non-uniform. Reconstructed aperiodic components in log–log space further highlighted this group-level steepening in LKO mice (Fig. 1F). These patterns were consistent across bilateral frontal and temporal channels, suggesting reorganization of cortical spectral dynamics during chronic epilepsy, though the changes in aperiodic exponent may be partly driven by undetected low-frequency epileptiform activity.

We next examined oscillatory (periodic) beta activity while controlling for aperiodic contributions. Subject-specific beta peaks were reconstructed using the individual central frequency for each animal, benchmarked against the canonical WT beta range. Beta central frequency was significantly lower in WT than in Epm2a−/− mice (Fig. 1G; n = 48 sessions; WT vs. LKO β = −3.08, 95% CI [−5.05, −1.04]; effect size = −1.08, 95% CI [−1.77, −0.37]).No interaction effect was observed. Beta power, adjusted for aperiodic components, was also significantly higher in LKO animals (Fig. 1H; 48 sessions; β = 0.15, 95% CI [0.03, 0.26]; effect size = 0.76, 95% CI [0.16, 1.35]), and this genotype difference was more pronounced in temporal cortex (groupLKO × temporal β = 0.05, 95% CI [0.001, 0.106]; effect size = 0.29, 95% CI [0.005, 0.55]). Additionally, aperiodic-adjusted delta power was elevated in LKO mice (**Fig. S2**; β = 0.18, 95% CI [0.02, 0.34]; effect size = 1.00, 95% CI [0.12, 1.92]), consistent with reports of cognitive impairment in this model.

Together, these findings reveal a coordinated shift in both aperiodic and periodic EEG features. Steeper aperiodic slopes, along with increased beta frequency and power are consistent with altered network-level spectral organization in Epm2a−/− mice at baseline, potentially reflecting progressive cortical adaptation to chronic epilepsy.

### 3.2 Isoflurane induced GABAergic modulation reveals genotype-dependent differences in spectral responsiveness in Epm2a−/− mice

Altered or immature GABAergic signaling is a well-established contributor to heightened seizure susceptibility, and seizures themselves can further disrupt GABA-A receptor function by modifying receptor subunits and altering inhibitory currents (Briggs and Galanopoulou., 2011). Although GABA-A receptor–acting anesthetics typically produce a steepening of the spectral slope, some modulators such as ketamine show inconsistent effects (Salvatore et al., 2024). Here we asked whether genotypic differences in baseline spectral organization shape the brain’s responsiveness to phasic GABAergic modulation, quantified through EEG-derived spectral features. Isoflurane was administered during bilateral 4-channel screw EEG recordings (Fig. 2A).

**Figure 2.**
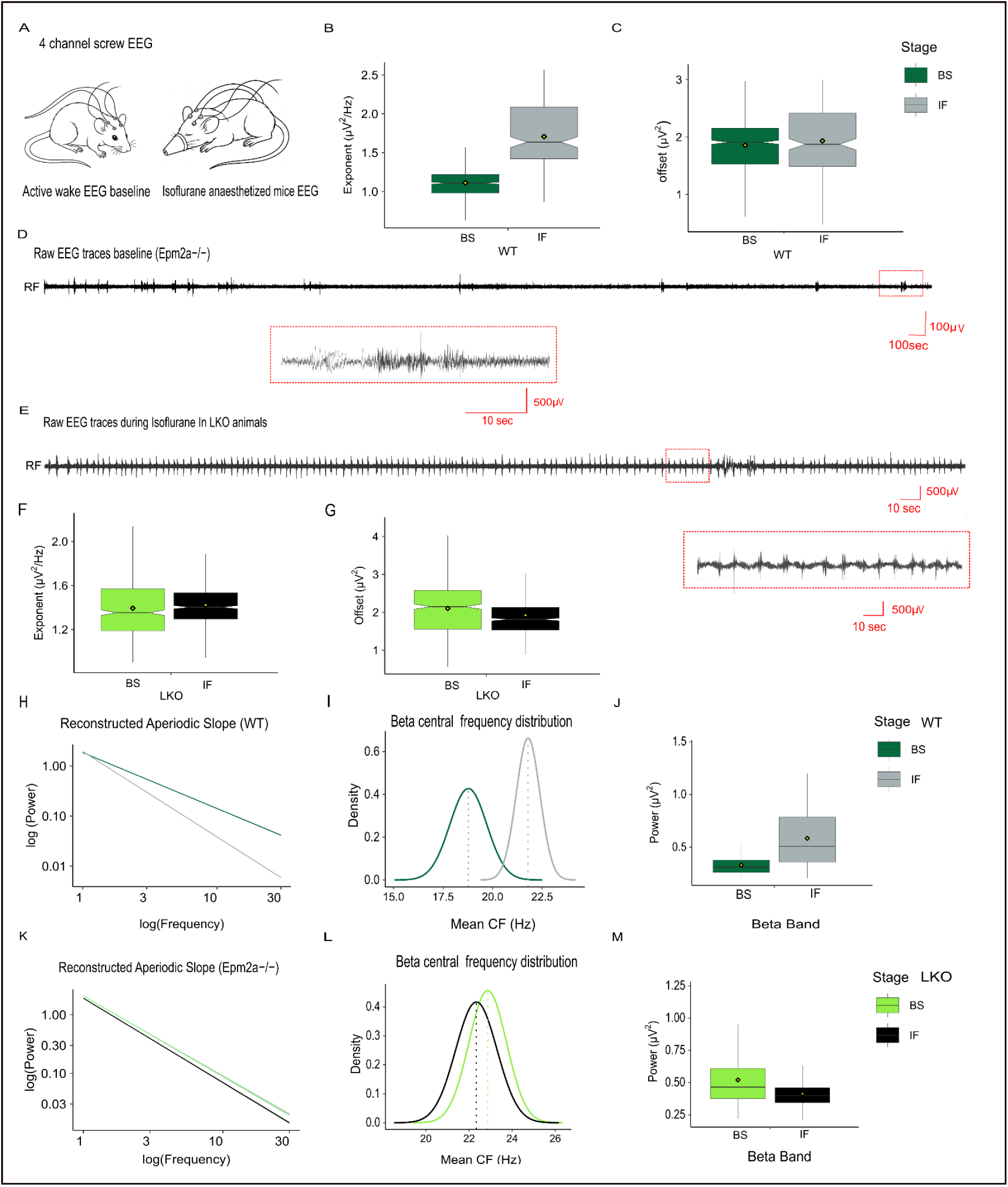
Isoflurane reveals genotype-dependent differences in aperiodic and oscillatory spectral responsiveness. **A.** Intracranial electrophysiological activity was recorded using screw EEG in C57BL/6 mice during wake active phase and during 1.5% Isoflurane infusion from two frontal, and two temporal screws, with the cerebellum as reference. **B.** Aperiodic exponent in C57BL/6 mice compared between baseline (Dark green) and 1.5% isoflurane (IF) (grey) conditions (scale: 100 μV/100 sec, zoomed at 500 μV/10 sec) **C.** Box plot illustrates no significant change in aperiodic offset after IF induction (grey) compared to baseline awake (Dark green) in C57BL/6 mice. **D.** Raw EEG traces from Epm2a−/− mice during task and seizure-free active wake state showing epileptic discharges resampled at 2000Hz. **E.** Raw EEG traces from Epm2a−/− mice during exposure to 1.5% isoflurane, indicating burst suppression patterns. **F.** Box plot comparing mean aperiodic exponent across all channels between baseline (light green) and 1.5% isoflurane (Black) in Epm2a−/− mice. **G.** Box plot comparing mean aperiodic offset across all channels between awake baseline (light green) and 1.5% isoflurane (Black) in Epm2a−/− mice. **H.** Reconstructed aperiodic component (after removing oscillatory peaks) showing steepening during isoflurane-induced pharmacological inhibition in WT mice. **I.** Comparison of canonical beta peaks with the deviation of the parameterized beta center frequency for the baseline (left, dark green) and isoflurane condition (right, grey) CF peak shift in canonical beta in C57BL/6 mice. **J.** Box plot illustrating aperiodic adjusted beta power comparison between baseline (dark green) and isoflurane (grey) epochs in WT (C57BL/6) mice. **K.** Reconstructed aperiodic component (after removing oscillatory peaks) showing steepening during isoflurane-induced pharmacological inhibition in C57BL/6 mice. **L.** Comparison of canonical beta peaks with the deviation of the parameterized beta center frequency between baseline (left, dark green) and isoflurane (right, grey), showing a CF peak shift in the beta band of Epm2a−/− mice **M.** Box plot illustrating a significant decrease in aperiodic adjusted beta power (µV²) in the canonical beta band during IF-induced cortical pharmacological inhibition in Epm2a−/− mice. Box plot represents median/IQR, notches are the 95% CI of the median, and the diamond is the mean.

In WT animals, isoflurane produced the expected broadband slowing of the EEG (1–30 Hz) and significantly steepening of the PSD, reflected by steeper aperiodic exponent (Fig. 2B; n = 8 animals, 32 sessions; β = 0.47, 95% CI [0.02, 0.94]; effect size = 2.51, 95% CI [0.11, 5.02]), with this effect being more pronounced in the temporal cortex ( n = 8, 32 sessions; β = 0.28, 95% CI [0.22, 0.34]; effect size = 1.49, 95% CI [1.18, 1.81]). Importantly, the aperiodic offset remained unchanged (Fig. 2C; n = 8, 32 sessions), However, interaction effect was observed in the temporal region ( β = 0.46, 95% CI [0.37, 0.57]; effect size = 1.35, 95% CI [1.05, 1.67]).This indicates that slope steepening reflected a redistribution of spectral power toward lower frequencies rather than a global power shift in frontal region (Fig. 2H).

Isoflurane also increased the aperiodic-adjusted beta central frequency (Fig. 2I; n = 8, 32 sessions; β = 3.07, 95% CI [1.52, 4.70]; effect size = 1.64, 95% CI [0.81, 2.55]). Elevated beta power, with this effect being more pronounced in the temporal cortex (Fig. 2J; n = 8, 32 sessions; β = 0.109, 95% CI [0.059, 0.159]; effect size = 0.787, 95% CI [0.42, 1.15]).

We next examined how Epm2a−/− mice responded to the same isoflurane concentration (1.5%) as WT animals. Representative EEG traces before and after induction are shown in Fig. 2D–E. Compared to WT, LKO animals displayed markedly different trends in both aperiodic and periodic components. Isoflurane decreased the aperiodic exponent relative to baseline in LKO mice (Fig. 2F; n = 8 animals, 32 sessions; β = −0.05, 95% CI [−0.09, −0.02]; effect size = −0.26, 95% CI [−0.42, −0.10]). No significance was observed with the region interaction model. Region-resolved analysis confirmed this divergent response: the exponent flattened only slightly relative to baseline ( n = 8 animals, 32 sessions; β = −0.04, 95% CI [−0.08, 0.00]; effect size = −0.19, 95% CI [−0.38, 0.00]), whereas the aperiodic offset dropped markedly (Fig. 2G; n = 8 animals, 32 sessions; β = −0.72, 95% CI [−0.84, −0.59]; effect size = −1.14, 95% CI [−1.34, −0.94]), indicating that anesthesia reduced broadband spectral power far more than it reshaped the 1/f slope; this offset reduction was attenuated in temporal relative to frontal region ( n = 8 animals, 32 sessions; Stage × temporal interaction β = 0.46, 95% CI [0.32, 0.60]; effect size = 0.73, 95% CI [0.50, 0.96]). Because the recordings under isoflurane showed burst-suppression patterns containing sharp transients, exponent estimates in these epochs should be interpreted with caution given the non-stationarity of burst-suppression activity. To assess whether burst-suppression-related waveform transients influenced the spectral estimates reported above, we applied SPRiNT-based decomposition over shorter 2-second epochs; similar state-dependent patterns were observed in both genotypes (Fig. S3).

After adjusting for aperiodic structure, isoflurane did not significantly change beta oscillatory frequency in LKO animals (Fig. 2L; n = 8 animals, 32 sessions) , but produced a true reduction in beta power (Fig. 2M; n = 8 animals, 32 sessions; β = −0.03, 95% CI [−0.05, −0.014]; effect size = −0.49, 95% CI [−0.69, −0.29]). Region-resolved analysis confirmed that the isoflurane-induced reduction in LKO beta power was significant in frontal cortex ( β = −0.035, 95% CI [−0.055, −0.015]; effect size = −0.28, 95% CI [−0.44, −0.12]) and significantly greater in temporal cortex (StageIF × temporal β = −0.063, 95% CI [−0.088, −0.037]; effect size = −0.50, 95% CI [−0.69, −0.29]). These combined changes suggest that baseline spectral differences and network-specific oscillatory mechanisms shape how the epileptic brain responds to isoflurane. This pharmacological manipulation also provided an orthogonal validation that exponent changes tracked physiological modulation of cortical state rather than disease alone.

### 3.3 Systematic aperiodic exponent variation across seizure phases is accompanied by pathological waveform contributions

Although seizures are traditionally linked to hyperexcitability, emerging cellular-level evidence shows that the E/I ratio does not rise uniformly; instead, it fluctuates dynamically across ictogenesis, with seizure initiation often driven by increased inhibition. To capture these evolving network states at the systems level, we again used aperiodic spectral parameters across pharmacologically induced seizures. Tonic–clonic (TC) seizures were triggered using the GABA-A receptor antagonist pentylenetetrazol (PTZ), enabling us to characterize preictal, ictal, and postictal epochs through detailed, epoch-by-epoch spectral parameterization (Fig. 3A–C). To assess the contribution of pathological waveform morphology to these exponent changes, we examined the correspondence between dominant waveform types present across seizure phases and their aperiodic spectral signatures. We hypothesized that the aperiodic exponent would vary systematically across seizure phases, and that pathological waveform morphology would emerge as a contributor to observed exponent changes during ictal epochs.

**Figure 3.**
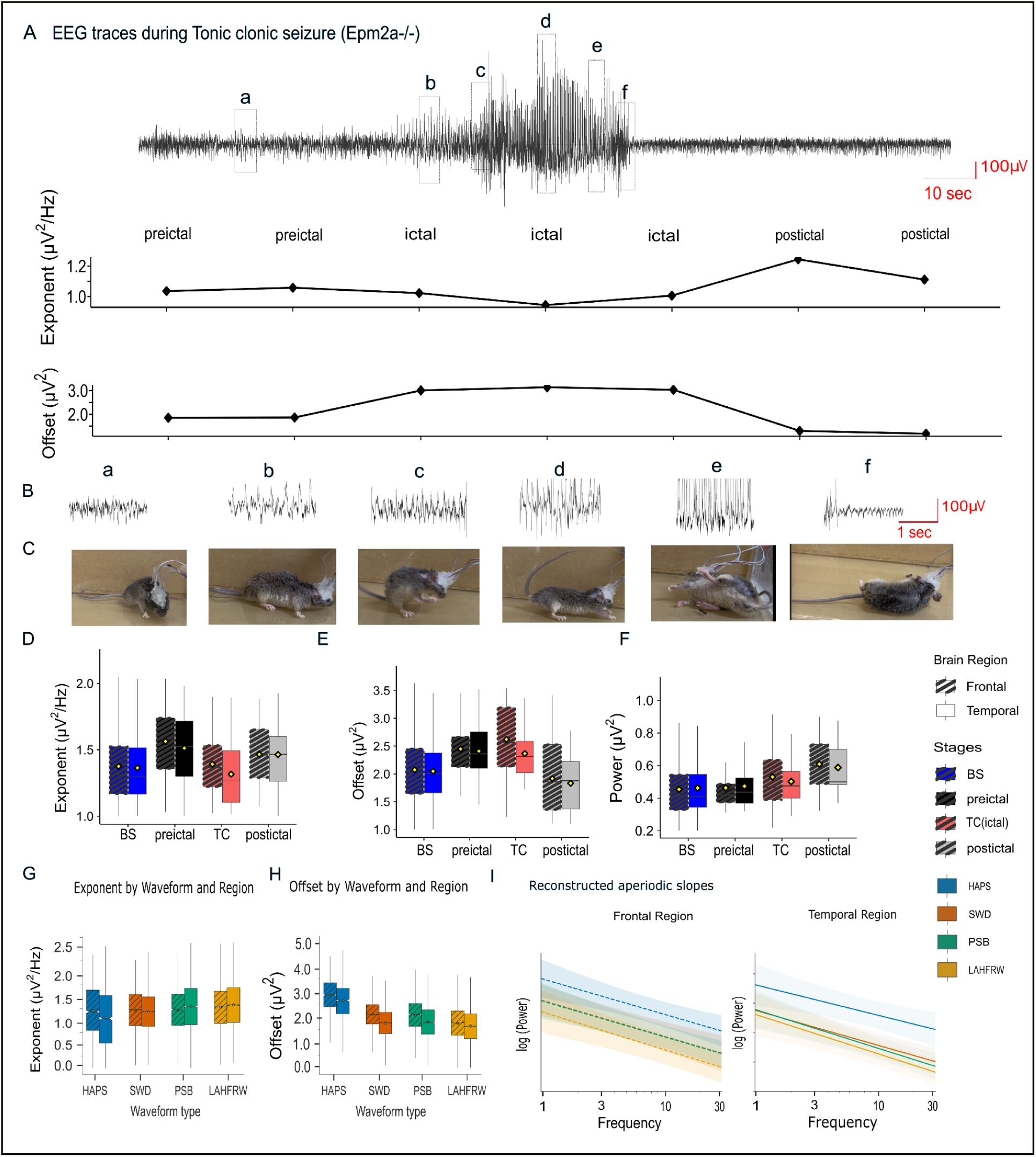
Aperiodic exponent varies systematically across seizure phases, with pathological waveform morphology as a contributing factor. **A.** Intracranial electrophysiological activity during tonic–clonic seizure in Epm2a−/− mice was recorded using screw EEG. Representative EEG trace from a single animal, the right temporal region is divided into preictal, ictal, and postictal stages following PTZ injection (55 mg/kg, i.p.) (scale: 100 μV/10 sec). A dynamic shift in the mean aperiodic exponent across seizure stages in this event is shown (middle), and a dynamic shift in the aperiodic offset, indicating changes in overall spectral power, is shown. Rectangular segments of the trace labeled a, b, c, d, e, f were zoomed to display aperiodic exponent, offset values, and behavioral phenotype with corresponding timestamps (seconds). **B.** Representative EEG features at corresponding timestamps (highlighted in the EEG trace with rectangles).**C.** Representative behavioral snapshots at different timepoints during the tonic–clonic seizure corresponding to EEG features with rectangles. **D.** Box plots showing changes in aperiodic exponent across tonic–clonic seizure stages (BS, preictal, ictal (TC), postictal) stages and brain regions (Frontal, temporal). **E.** The box plot illustrates dynamic offset across baseline, preictal, ictal and postictal across brain regions (Frontal, Temporal) **F.** Statistical illustration of aperiodic-adjusted beta oscillatory power across stages baseline, preictal, ictal and postictal across brain regions (Frontal, Temporal). **G.** Box plots of aperiodic exponent across pathological waveform types (HAPS, SWD, PSB, LAHFRW) and brain regions; relative to HAPS, all other waveform types showed significantly higher (steeper) exponents. **H.** Box plots of aperiodic offset across the same waveform types and regions; relative to HAPS, all other waveform types showed significantly lower offsets. **I.** Reconstructed aperiodic slopes in log–log space for the frontal (left) and temporal (right) regions, showing the flatter slope and higher offset associated with HAPS compared with SWD, PSB, and LAHFRW. Box plot represents median/IQR, notches are the 95% CI of the median, and the diamond is the mean. Baseline frontal (blue, patterned), Baseline temporal (blue, plain), preictal frontal (black, patterned), preictal temporal (black, plain), ictal frontal (red, patterned), ictal temporal (red, plain), post-ictal frontal (grey, patterned), ictal temporal (grey, plain).Waveforms represented High Amplitude Poly Spikes (HAPS), SWD (Spike Wave Discharge), Poly Spike Burst (PSB), Low Amplitude High Frequency Rhythmic Waves (LAHFRW).

The aperiodic exponent, reflecting the 1/f spectral slope, varied systematically across seizure phases (Fig. 3D). Relative to baseline, the exponent steepened significantly during the preictal period (n = 8 animals; β = 0.20, 95% CI [0.14, 0.27]; effect size = 0.81, 95% CI [0.55, 1.07]), indicating a steeper slope across frontal and temporal cortices. During the ictal (TC) phase the exponent showed a marked flattening relative to the preictal ( n = 8, 24 sessions; β = −0.20, 95% CI [−0.28, −0.12]; effect size = −0.79, 95% CI [−1.09, −0.48]). The exponent also showed significant flattening during TC from baseline in temporal region (vs baseline: n = 8, 24 sessions; β = −0.11, 95% CI [−0.19, −0.03]; effect size = −0.43, 95% CI [−0.74, −0.11]). Following seizure termination, the exponent again exceeded baseline (vs baseline: β = 0.15, 95% CI [0.04, 0.25]; effect size = 0.58, 95% CI [0.18, 0.98]), reflecting a rebound in spectral slope consistent with postictal suppression.

The aperiodic offset, indexing broadband power, also showed phase-specific modulation (Fig. 3E). Relative to baseline, the offset increased significantly during the preictal period (n = 8 animals; β = 0.41, 95% CI [0.27, 0.55]; effect size = 0.74, 95% CI [0.48, 1.00]) and rose further during the ictal (TC) phase (β = 0.93, 95% CI [0.78, 1.09]; effect size = 1.69, 95% CI [1.40, 1.97]), indicating a marked increase in broadband spectral power at ictus. Postictally, the offset returned toward baseline and no longer differed significantly from it; however, relative to the elevated preictal level, the postictal offset dropped markedly (n = 8, 24 sessions; β = −0.56, 95% CI [−0.79, −0.32]; effect size = −1.01, 95% CI [−1.43, −0.59]), consistent with a strong broadband-power collapse during postictal suppression. No effect of interaction observed.

Periodic oscillatory EEG features also tracked seizure-related state changes (Fig. 3F). Relative to preictal levels, beta-band power increased during the ictal (TC) phase (β = 0.14, 95% CI [0.005, 0.27]; effect size = 0.74, 95% CI [0.03, 1.47]) and remained elevated after seizure termination, with postictal beta power significantly higher than preictal (β = 0.25, 95% CI [0.07, 0.42]; effect size = 1.33, 95% CI [0.39, 2.28]); no significant regional interactions were observed.

Elevations in aperiodic-adjusted power during the ictal (TC) phase relative to baseline were band-specific: sigma and alpha power increased significantly, whereas delta, theta, and beta power did not differ significantly from baseline as well as from preictal (**Fig. S4**), indicating selective reorganization of mid-frequency rhythms during ictogenesis. Aperiodic-adjusted beta central frequency increased significantly during the ictal phase relative to preictal levels (**Fig. S7**).

To directly assess the contribution of pathological waveform morphology to the seizure-phase exponent changes described above, we fit a separate model relating the aperiodic exponent to waveform type (high amplitude poly spike (HAPS), spike-wave discharges (SWD), polyspike bursts (PSB), and large-amplitude high-frequency rhythmic waveforms (LAHFRW) in temporal cortex (Fig. 3G). Relative to HAPS, all three other waveform types were associated with significantly higher aperiodic exponents: LAHFRW (β = 0.25, 95% CI [0.23, 0.28]; effect size = 0.55, 95% CI [0.49, 0.61]), PSB (β = 0.20, 95% CI [0.18, 0.23]; effect size = 0.45, 95% CI [0.38, 0.51]), and SWD (β = 0.09, 95% CI [0.06, 0.12]; effect size = 0.20, 95% CI [0.14, 0.27]), pronounced in temporal region. Thus HAPS, the waveform dominating high-grade ictal activity, carried the lowest (flattest) aperiodic exponent, consistent with the ictal exponent flattening reported above.

To assess how waveform morphology related to broadband power, we modeled the aperiodic offset as a function of waveform type (Fig. 3H). Relative to HAPS, all three other waveform types were associated with significantly lower offsets in temporal region: LAHFRW (β = −1.36, 95% CI [−1.39, −1.33]; effect size = −2.17, 95% CI [−2.22, −2.12]), PSB (β = −1.08, 95% CI [−1.12, −1.05]; effect size = −1.73, 95% CI [−1.78, −1.67]), and SWD (β = −1.07, 95% CI [−1.10, −1.03]; effect size = −1.70, 95% CI [−1.76, −1.65]). Reconstructed aperiodic slopes in log–log space illustrate this waveform-specific spectral signature across frontal and temporal regions (Fig. 3I). Thus HAPS, the waveform dominating high-grade ictal activity, carried the highest broadband offset, consistent with the elevated ictal offset reported above.

Together, these results reveal an aperiodic spectral dynamics: a steeper-slope preictal period, a transition to a flatter slope during the ictal phase, and a strong slope rebound during postictal suppression. However, these phase-dependent exponent changes are accompanied by systematic variation in pathological waveform morphology, and the relative contributions of true network-state reorganization versus waveform-shape-driven spectral distortion cannot be fully disentangled using spectral parameterization alone.

### 3.4 Aperiodic exponent scales with seizure grade and reflects waveform-shape contributions

Our earlier analyses revealed systematic exponent variation across tonic–clonic (TC) seizure phases. We next examined whether lower-intensity seizures, particularly those lacking a clear preictal stage are driven primarily by distinct spectral dynamics, and whether aperiodic parameters can reliably differentiate seizure severity. To address this, we analyzed EEG epochs across multiple seizure grades. Based on behavioral features, we hypothesized that grade-2 seizures, which frequently include spike–wave discharges, would produce steeper the aperiodic slope, whereas grade-4 seizures, characterized by stronger motor involvement, greater exponent flattening.

The aperiodic exponent was sensitive to seizure grade, offering a continuous, noninvasive marker of cortical network state(Fig. 4). Representative traces from LKO animals illustrate grade-2 seizures marked by spike–wave discharges and polyspikes, and grade-4 seizures characterized by higher-amplitude ictal spikes (Fig. 4A).

**Figure 4.**
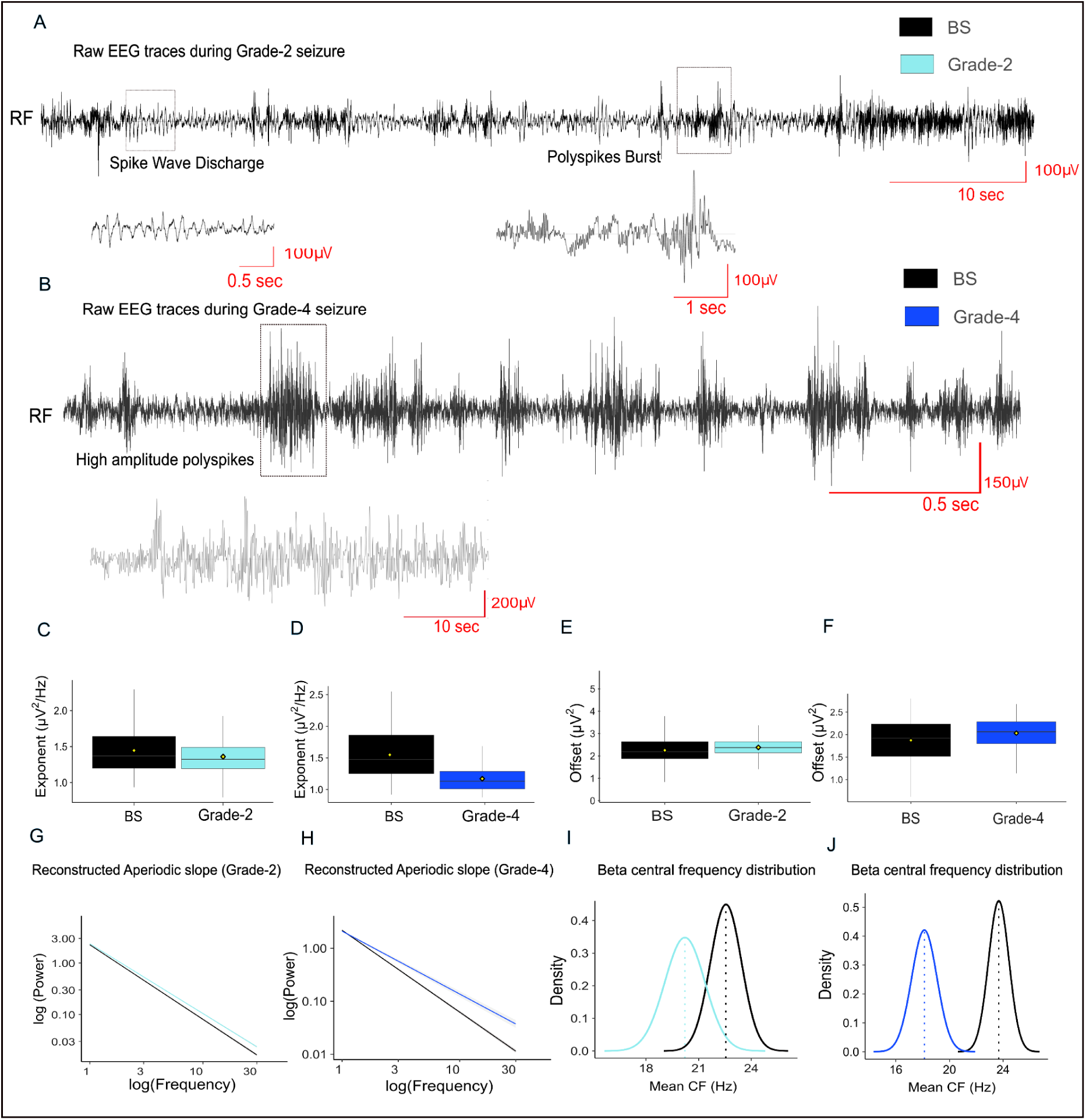
Aperiodic exponent scales with seizure grade alongside graded shifts in dominant waveform morphology. **A.** Intracranial electrophysiological activity during low grade seizure in Epm2a−/− mice was recorded using screw EEG. Representative trace from the right frontal brain region for epochs of low grade seizure stages induced after PTZ injection (55mg/kg, i.p, n = 8, session=32), scale 100μV/10sec, zoomed at 100μV/1sec, Spike-wave discharges and Polyspikes burst are the dominant waveform types in this grade. **B.** Representative trace from the right frontal brain region for epochs of higher grade seizure stages induced after PTZ injection ( n = 4, session=26), scale 200μV/10sec, zoomed at 150μV/0.5sec, High-amplitude poly spikes are the dominant waveform type in this grade **C.** Lower aperiodic exponent (indicating flattening) during grade 2 seizure (sky blue) compared to baseline (black). **D.** Lowerer aperiodic exponent (indicating flattening) during grade 4 seizure (dark blue) compared to baseline and grade 2 seizure (black). **E.** Moderate change in aperiodic offset during grade 2 seizure compared to baseline. **F.** Further elevation in aperiodic offset during grade 4 seizure (dark blue) compared to baseline (black). **G.** Reconstructed aperiodic slope (log-log) indicating flattening during grade-2 seizure dominating 30s epochs. **H.** Reconstructed aperiodic slope (log-log) indicating flattening during grade-4 seizure dominating 30s epochs. **I.** Periodic Central Frequency (12-30Hz) peak in beta frequency band changes across stages baseline, grade-2 and **J.** grade-4 in *Epm2a−/− in* throughout all (LF, RF, LT, RT) brain regions. Gaussian density plots depict the distribution of CF peaks across epochs within each condition. Box plot represents median/IQR, notches are the 95% CI of the median, and the diamond is the mean. Screw EEG channel LF (Left Frontal), RF (Right Frontal), LT (Left Temporal), RT (Right Temporal).

During grade-2 seizures, the aperiodic exponent significantly flatters relative to baseline (Fig. 4C; n = 8, 32 sessions; β = −0.13, 95% CI [−0.16, −0.09]; effect size = −0.53, 95% CI [−0.68, −0.40]), an effect that was significantly attenuated in temporal relative to frontal cortex (Grade-2 × temporal β = 0.08, 95% CI [0.04, 0.13]; effect size = 0.36, 95% CI [0.17, 0.56]). Notably, grade-2 seizures were dominated by spike-wave discharges and polyspike-burst waveform types, whose spectral profiles are consistent with steeper aperiodic slopes compared to TC dominated HAPS, but flatter compared to BS.

To test whether these spectral shifts scale with seizure intensity, we next examined grade-4 seizures. In these higher-grade epochs, the aperiodic exponent showed a more pronounced flattening than in grade-2 seizures (Fig. 4D; n = 4, 26 sessions; β = −0.20, 95% CI [−0.34, −0.05]; effect size = −0.68, 95% CI [−1.17, −0.18]).No interaction effect was observed. Grade-4 seizures were characterized by high-amplitude ictal spikes, which similarly carry spectral signatures consistent with flatter slopes, indicating that the graded exponent decrease across seizure severity is paralleled by a graded shift in the dominant pathological waveform type.

Aperiodic offset values also scaled with seizure intensity. During grade-2 seizures, the offset increased significantly relative to baseline (Fig. 4E; n = 8, 32 sessions; β = 0.17, 95% CI [0.11, 0.23]; effect size = 0.40, 95% CI [0.26, 0.55]), indicating greater overall spectral power. No interaction observed. Although grade-4 seizures also showed a numerical increase in offset relative to baseline, this effect did not reach significance (Fig. 4F; n = 4, 26 sessions). However, within grade-4 seizures, temporal regions showed a significantly lower offset than frontal cortex (**Fig. S5**; n = 4, 26 sessions; region β = −0.43, 95% CI [−0.54, −0.31]), effect size = −0.01, 95% CI [−0.65, 0.64]), suggesting region-specific differences in high-grade ictal recruitment.

Consistent with these aperiodic effects, aperiodic slope flattening was evident in both seizure grades, with grade-4 seizures exhibiting the most pronounced flattening (Fig. 4G–H), and more prominent high-amplitude ictal waveforms. The aperiodic exponent and offset appeared to be better temporally resolved markers of seizure severity since we did not observe significant change in Hurst exponent among the stages and grades of seizure (**Fig.S6).**

Together, these results show that both grade-2 and grade-4 seizures are marked by flatter aperiodic slopes, with progressively lower exponents and elevated offsets as seizure intensity increases. However, the graded exponent decrease across seizure grades is systematically paralleled by a graded shift in dominant pathological waveform morphology, indicating that waveform-shape-driven spectral distortion contributes to and cannot be fully dissociated from, the observed grade-dependent exponent changes using spectral parameterization alone.

## 4. Discussion

Chronic epilepsy involves progressive, brain-wide alterations in network organization, yet clinical recordings are frequently confounded by anti-epileptic medications that obscure intrinsic homeostatic adaptations. Using a chronic epileptic knockout model, we examined drug-free network dynamics and show how the aperiodic exponent of the EEG power spectrum provides a sensitive, temporally resolved descriptor of cortical network state across seizure stages and severities. Here, we examined how the aperiodic EEG spectrum evolves across chronic epilepsy, pharmacological modulation, and seizure progression in a medication-free genetic model. Across these conditions, aperiodic dynamics varied systematically with brain state, suggesting that the physiological interpretation of the exponent is itself state-dependent.

Baseline recordings revealed steeper aperiodic slopes in Epm2a−/− mice, consistent with altered cortical network organization and possibly greater inhibitory tone. Although these mice show early-life reductions in cortical GABAergic neurons (Ortolano et al., 2014), inhibition can remain preserved or even increase in epilepsy (Cope et al., 2009; Müller et al., 2020). Importantly, excessive tonic inhibition may paradoxically increase seizure vulnerability by narrowing the dynamic range of inhibitory control and creating a brittle network prone to collapse during excitatory perturbations (Mathew et al., 2012; Mann & Mody, 2008). This interpretation aligns with molecular evidence for altered inhibitory homeostasis in Epm2a−/− mice (Ganesh et al., 2005) and with human recordings showing altered aperiodic slopes during interictal periods (Kopf et al., 2024; Kozma et al., 2024). Importantly, the steeper exponent in LKO animals persisted after excluding epochs containing interictal epileptic discharges, suggesting that this baseline spectral difference reflects chronic network-level reorganization rather than being driven by the presence of pathological waveforms. This establishes the resting aperiodic exponent as a robust marker of sustained network state change in chronic epilepsy, independent of acute waveform contamination.

Isoflurane anesthesia produced robust slope steepening, consistent with prior reports that GABAergic anesthetic states alter aperiodic spectral structure (Brown et al., 2010; Purdon et al., 2015), with distinct genotype-dependent patterns. Wild-type animals showed pronounced increases in beta power and frequency, whereas Epm2a−/− mice displayed altered modulation, likely reflecting disrupted GABA-A receptor signaling or interneuron recruitment (Jensen et al., 2005; Bieda et al., 2009). Because beta synchronization arises from coordinated interneuronal activity, baseline spectral organization and GABAergic responsiveness are differentially altered across genotypes in chronic epilepsy. However, it is worth noting that all recordings under isoflurane showed burst-suppression patterns with sharp transients in both genotypes; exponent estimates in these epochs may therefore partially reflect non-stationarity of burst-suppression activity rather than smooth spectral reorganization, and should be interpreted accordingly. Importantly, identical GABAergic manipulation produced different spectral consequences because the underlying network state differed.

During PTZ-induced seizures, the aperiodic exponent varied systematically across seizure phases: steepening during the preictal period, flattening during the ictal phase, and steepening again during postictal suppression. Importantly, these transitions occurred within the same animals over short time scales, suggesting that aperiodic dynamics capture evolving network state rather than fixed disease characteristics. These observations are broadly consistent with a two-step model of ictogenesis in which inhibition first increases to stabilize the network, followed by a transition to sustained excitation that drives seizure propagation (Žiburkus et al., 2013; Rich et al., 2020; Lőrincz et al., 2024). We emphasize that the aperiodic exponent provides a spectral index of these transitions rather than a direct measure of synaptic E/I ratios. The exponent also scaled with seizure grade, flattening more strongly during high-intensity seizures and enabling fine-grained differentiation of seizure severity beyond behavioral scoring alone.

Our waveform analyses further demonstrate that the physiological interpretation of the aperiodic exponent is itself state-dependent. During resting wakefulness, isoflurane anesthesia, and peri-ictal periods, exponent changes behaved in a manner broadly consistent with evolving cortical network states. During sustained ictal activity, however, pathological waveform morphology increasingly dominated spectral slope estimates. This observation aligns with recent work showing that nonsinusoidal waveform features and periodic structure can distort estimates of the aperiodic component obtained through spectral parameterization methods (Donoghue et al., 2020; Gerster et al., 2022; Aggarwal and Ray, 2025; Heidiri et al., 2025). Our findings therefore suggest that aperiodic dynamics in epilepsy arise from a combination of genuine network-state reorganization and waveform-shape-driven spectral distortion, with their relative contributions depending on the underlying physiological state. Importantly, this does not diminish the utility of spectral parameterization as a biomarker, but instead defines the conditions under which its interpretation is likely to be most informative.

Our analyses relied on low-channel EEG and a single genetic model of chronic epilepsy, constraining spatial resolution and the generalizability of absolute quantitative values. Future work applying this framework to high-density EEG, intracranial recordings, and human datasets will be essential for testing its generality across species and disease contexts. Normative mapping of interictal human iEEG and MEG offers one such route, though notably the aperiodic exponent in isolation did not distinguish surgical outcome in that setting, whereas combining periodic and aperiodic abnormalities did (Kozma et al., 2024) underscoring that the exponent is best interpreted alongside, rather than in place of, periodic spectral features. Future analyses using complementary approaches such as IRASA or knee-parameterized models could further test the robustness of these slope estimates. Although baseline sessions were curated as wake, we did not include physiological markers of vigilance (e.g., EMG/pupil), and residual arousal differences could influence the aperiodic exponent. In addition, PTZ-induced seizures provide controlled access to pre-ictal, ictal, and post-ictal states but do not capture the full heterogeneity of spontaneous epilepsy. Furthermore, directly disentangling the contributions of true network-state reorganization from waveform-shape-driven spectral distortion will require concurrent intracellular or laminar recordings that provide ground-truth measures of synaptic excitation and inhibition alongside surface EEG. At the same time, the within-animal design, absence of AED confounds, and combination of chronic genetic epilepsy with pharmacological manipulation provide a unique opportunity to dissociate network state from medication effects.

More broadly, these findings argue that aperiodic activity should be interpreted as a dynamic descriptor of network organization whose physiological meaning depends on brain state. Although developed here in a chronic epilepsy model, this framework has broader implications for applying spectral parameterization to sleep, anesthesia, neurodegeneration, neuropsychiatric disorders, and other disease- or treatment-altered recordings.

## ACKNOWLEDGEMENTS

We are grateful to Dr. Rashmi Parihar and staff of the Central Experimental Animal Facility, IIT Kanpur, for their support in providing and maintaining animals . We thank Ananya Avasthy for her assistance with citation referencing. In Fig1&2 mice schematic illustrations were generated with assistance from an AI-based image generation tool (ChatGPT, OpenAI). All figure concepts, scientific content, and final modifications were designed, reviewed, and validated by the authors to ensure accuracy and compliance with PNAS Nexus.

## Artificial Intelligence (AI) Disclosure

No AI tool was used to design the study, acquire or analyze data, generate or interpret results, or produce any data-derived figure. All figure concepts, scientific content, and final modifications were designed, reviewed, and validated by the authors. Generative AI tools were used for language editing and for checking internal consistency of citations and cross-references in the manuscript text. Schematic illustrations of mice in Figures 1 and 2 were generated with assistance from an AI-based image generation tool (ChatGPT-5.1, OpenAI, Claude 4.8 Opus).

## AUTHOR CONTRIBUTIONS

Garima Chauhan (GC) and Arjun Ramakrishnan (AR) conceived and designed the study. GC, Kshitij Kumar (KK), and Deepti Chugh (DC) performed the animal surgeries and data acquisition. KK conducted the formal analysis of neural recordings and generated the visualizations. GC and KK wrote the initial draft of the manuscript. Subramaniam Ganesh (SG) developed the transgenic mouse model used in this study. All authors contributed to manuscript revision and approved the final version.

