## Supplementary File for "State-dependent aperiodic EEG dynamics track cortical network reorganization in chronic epilepsy"

**S1. Effect of interictal epileptiform discharges on aperiodic parameters**

**Cited in results 3.1**

**Time-resolved spectral parameterization (SPRiNT) analysis results:**

Because interictal epileptiform discharges (IEDs) were present in the EEG traces of LKO mice, we applied SPRiNT (Spectral Parameterization Resolved in Time; Wilson et al., 2022) to minimize the influence of these transient waveforms on Fourier-based spectral decomposition. IED-containing windows constituted 4.8% of the LKO baseline, and the exponent recomputed from the remaining IED-free windows (95.2%) remained steeper in LKO than in WT mice. IED occurrence did not significantly affect the aperiodic exponent within the genotype. The number of IEDs differed significantly between LKO and WT mice. The aperiodic offset did not differ significantly by genotype or IED status.

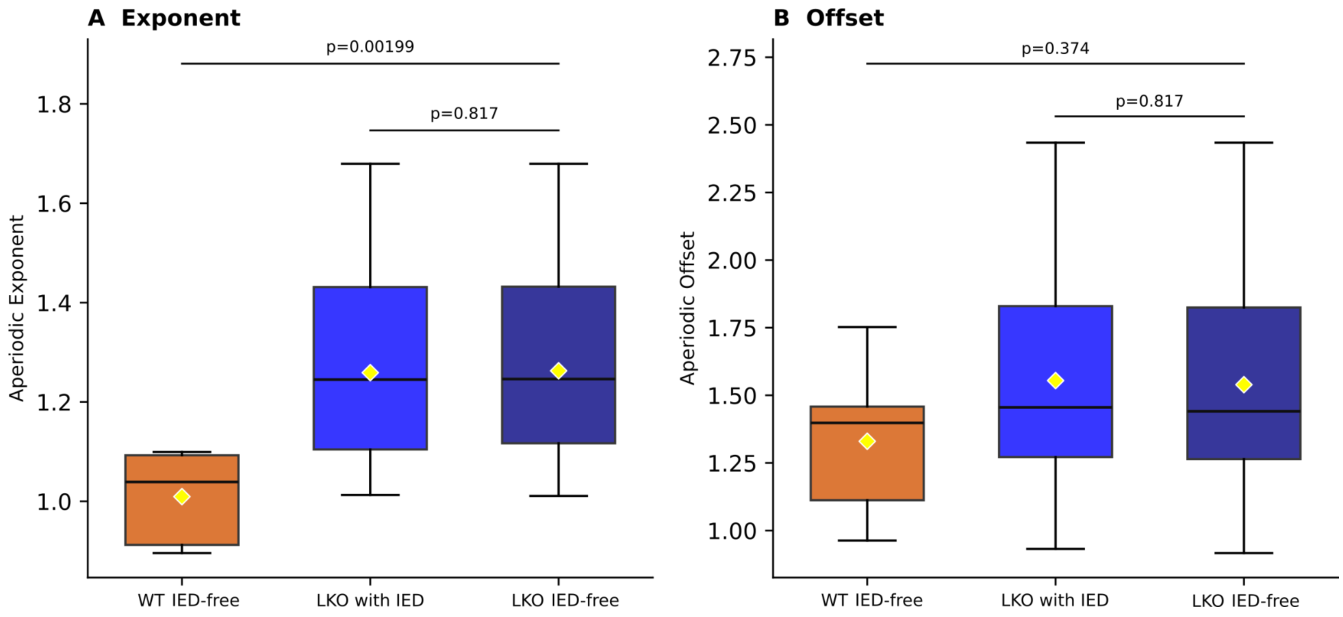


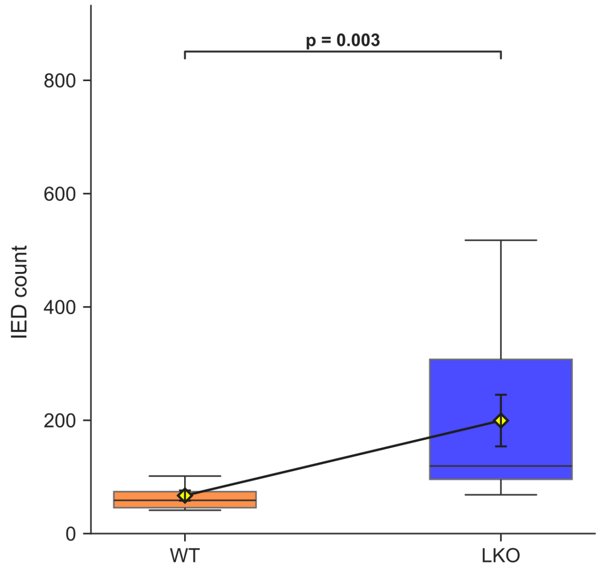

**Fig. S1.** Aperiodic exponent and offset in WT and LKO mice, with and without interictal epileptiform discharges (IEDs). (A) Aperiodic exponent was significantly steeper in LKO mice than in WT mice, irrespective of IED presence (LKO IED-free vs. WT IED-free, P = 0.002); exponent did not differ significantly between LKO recordings with and without IEDs (P = 0.82). (B) Aperiodic offset did not differ significantly by genotype (P = 0.37) or IED status (P = 0.82). Boxes show median and interquartile range; whiskers show data range; diamonds indicate group means. The Mann-Whitney U test is used to compare the distributions of two groups.

**S2. Aperiodic-adjusted delta-band power**

**Cited in: Results 3.1. FOOOF spectral parameterization.**

**FOOOF analysis results:**

Aperiodic-adjusted delta power was elevated in LKO mice, consistent with reports of cognitive impairment in this model.

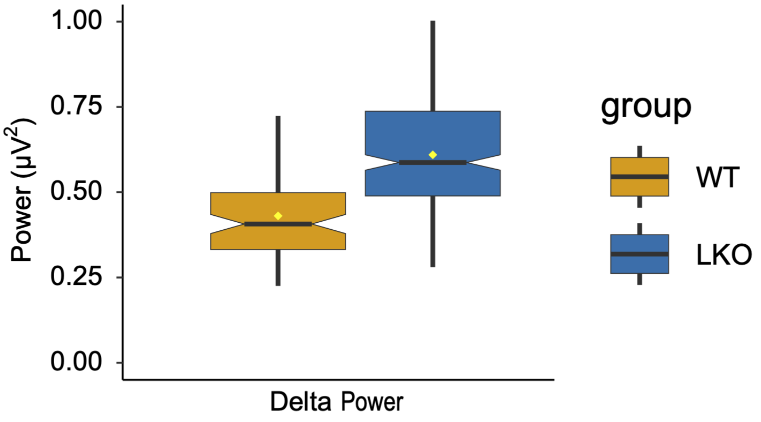


Fig. S2. Aperiodic-adjusted delta-band power in WT and LKO mice, estimated using FOOOF (fitting oscillations & one-over-f) spectral parameterization. Additionally, aperiodic-adjusted delta power was elevated in LKO mice (Fig. S2; β = 0.18, 95% CI [0.02, 0.34]; effect size = 1.00, 95% CI [0.12, 1.92]), consistent with reports of cognitive impairment in this model. No significant effect of interaction was observed. Box shows median and interquartile range (notch denotes the 95% confidence interval around the median); whiskers show data range; diamonds indicate group means.

**S3. Effect of isoflurane on the aperiodic exponent, controlling for burst-suppression transients**

**Cited in: Results 3.2. Time-resolved spectral parameterization (SPRiNT), 2-s epochs.**

To assess whether burst-suppression-related waveform transients influenced the spectral estimates reported above, we applied SPRiNT-based decomposition over shorter 2-second epochs. Isoflurane (IF) exposure at 1.5% produced a significantly greater steepening of the aperiodic exponent in WT mice than in LKO mice.

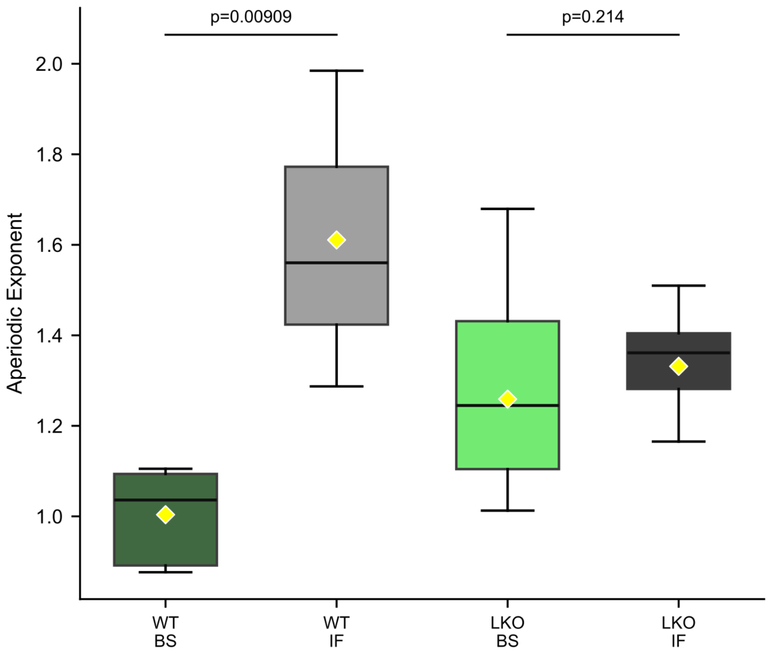


**Fig. S3.** Effect of isoflurane (IF) anesthesia on the aperiodic exponent in WT and LKO mice. SPRiNT-based decomposition was applied over 2-s epochs to control for burst-suppression-related waveform transients. IF exposure (1.5%) produced a significantly greater steepening of the aperiodic exponent in WT mice (baseline vs. IF, P = 0.009) than in LKO mice (baseline vs. IF, P = 0.21). BS, (baseline). Boxes show median and interquartile range; whiskers show data range; diamonds indicate group means.The Mann-Whitney U test is used to compare the distributions of two groups.

**S4. Aperiodic-adjusted band power across seizure stages**

**Cited in: Results 3.3.**

Aperiodic-adjusted power was compared across seizure stages, including the tonic–clonic (TC) stage and surrounding baseline, preictal, and postictal periods, for the alpha, beta, delta, sigma, and theta bands.

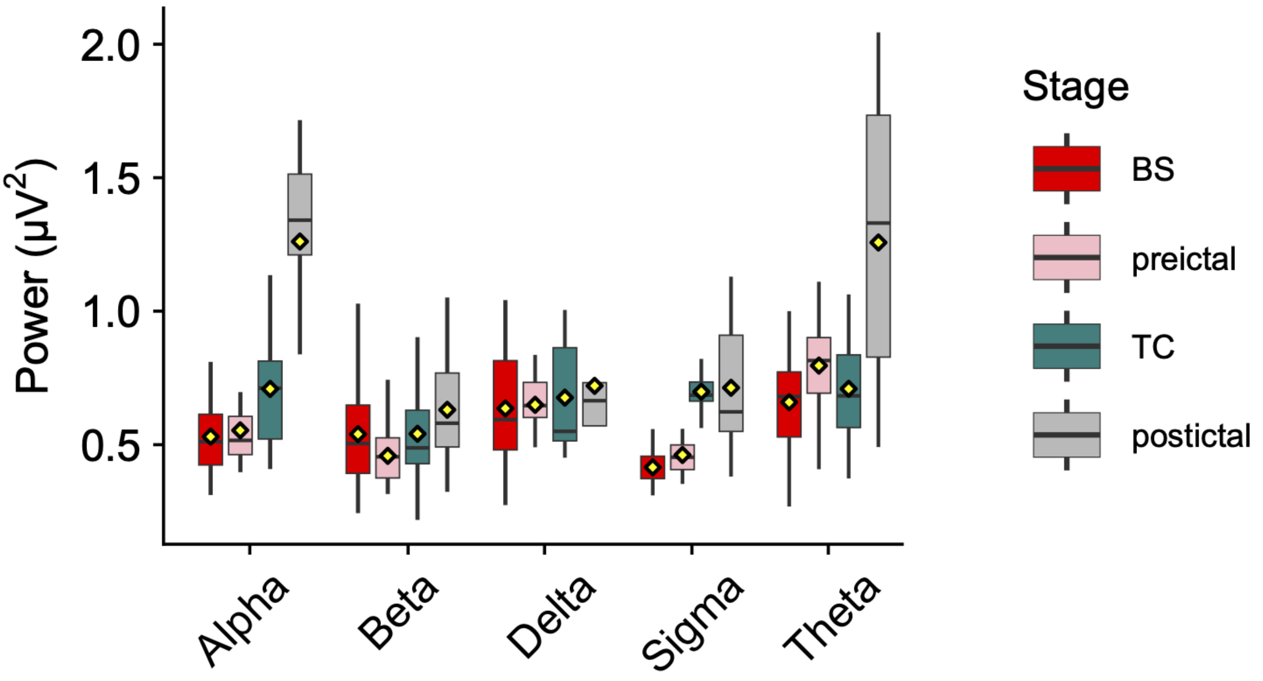


**Fig. S4.** Elevations in aperiodic-adjusted power during the ictal (TC) phase relative to baseline were band-specific: sigma (β = 0.29, 95% CI [0.23, 0.34]; effect size = 2.68, 95% CI [2.04, 3.37]) and alpha power (β = 0.18, 95% CI [0.10, 0.26]; effect size = 0.83, 95% CI [0.45, 1.20]) increased significantly from baseline, whereas delta, theta, and beta power did not differ significantly from baseline (Fig. S4), indicating selective reorganization of mid-frequency rhythms during ictogenesis. Relative to preictal levels, sigma power was significantly elevated during the ictal (TC) phase (β = 0.24, 95% CI [0.17, 0.31]; effect size = 2.24, 95% CI [1.55, 2.95]), as was alpha power (β = 0.15, 95% CI [0.008, 0.29]; effect size = 0.69, 95% CI [0.04, 1.36]); delta, theta, and beta power did not differ significantly from preictal during the ictal phase. The most consistent band-power changes were postictal: alpha (β = 0.70, 95% CI [0.55, 0.86]; effect size = 3.24, 95% CI [2.45, 4.04]), theta (β = 0.47, 95% CI [0.33, 0.61]; effect size = 2.10, 95% CI [1.44, 2.77]), beta (β = 0.18, 95% CI [0.07, 0.30]; effect size = 0.97, 95% CI [0.34, 1.61]), and sigma (β = 0.25, 95% CI [0.17, 0.34]; effect size = 2.36, 95% CI [1.51, 3.21]) power all increased significantly relative to preictal, whereas delta power showed no significant modulation at any phase. Boxes show median and interquartile range; whiskers show data range; diamonds indicate group means.

**S5. Aperiodic-adjusted beta-band activity during grade-2 and grade-4 seizures**

**Cited in: Results 3.4.**

**Beta-band activity:** We examined aperiodic-adjusted beta-band activity during moderate (grade-2) and severe (grade-4) seizures. In grade-2 seizures, beta central frequency (CF) was significantly reduced relative to baseline (main Fig. 4I; n = 8 mice, 32 sessions; β = −1.67, 95% CI [−2.07, −1.27]; effect size = −0.59, 95% CI [−0.64, 0.32]), accompanied by a significant reduction in beta power (n = 8 mice, 32 sessions; β = −0.04, 95% CI [−0.06, −0.02]; effect size = −0.27, 95% CI [−0.40, −0.14]), with no region-specific differences observed. In grade-4 seizures, beta CF was also significantly reduced relative to baseline (n = 4 mice, 26 sessions; 95% CI [−3.5, −0.75]; effect size = −0.78, 95% CI [−1.31, −0.27]), while beta power trended lower but did not reach statistical significance.

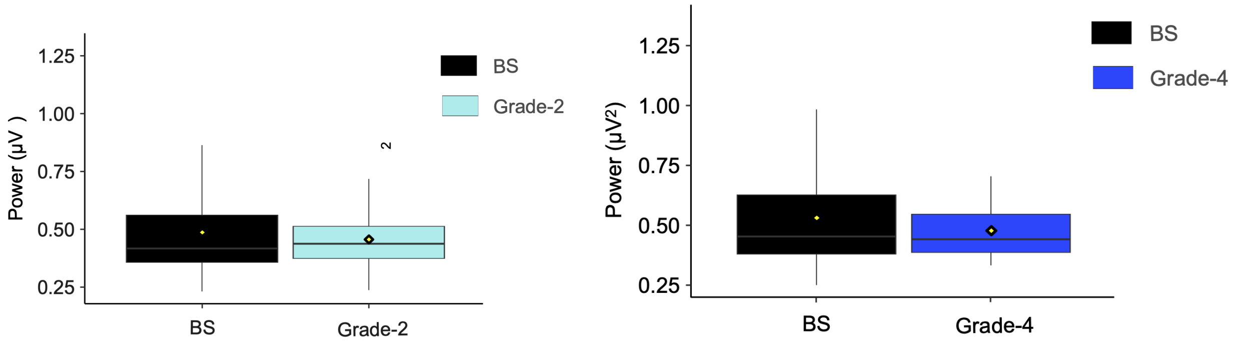


**Fig. S5.** Aperiodic-adjusted beta-band power during grade-2 and grade-4 seizures relative to baseline (BS). (Left) Beta power during grade-2 seizures compared with baseline (n = 8 mice, 32 sessions). (Right) Beta power during grade-4 seizures compared with baseline (n = 4 mice, 26 sessions). Boxes show median and interquartile range; whiskers show data range; diamonds indicate group means.

**S6 Hurst exponent Seizure Grade wise Comparison**

**Cited in: Results 3.4.**

As a complementary, model-free measure of long-range temporal correlations in the EEG signal, we computed the Hurst exponent (H) across seizure grades and stages. H did not differ significantly between the preictal, grade-2, grade-4, tonic–clonic (TC), and postictal periods, and remained above 0.5 across all stages.

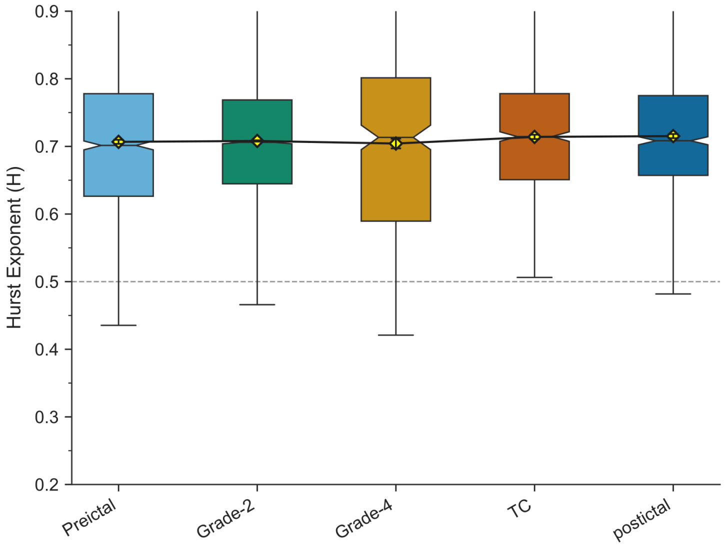


**Fig. S6.** Fig. S6. Hurst exponent (H) across seizure grades and stages (preictal, grade-2, grade-4, tonic–clonic (TC), and postictal). H did not differ significantly between grades or stages. Boxes show median and interquartile range (notches denote the 95% confidence interval around the median); whiskers show data range; diamonds indicate group means.

**S7 Phasic change in Aperiodic exponent and offset with Beta central Frequency**

**Cited in: Results 3.3.**

Aperiodic-adjusted beta central frequency increased significantly during the ictal phase relative to preictal levels (n = 8, 24 sessions; β = 2.28, 95% CI [0.96, 3.70]; effect size = 0.74, 95% CI [0.31, 1.20]), with no significant regional differences and no significant change at other phases.

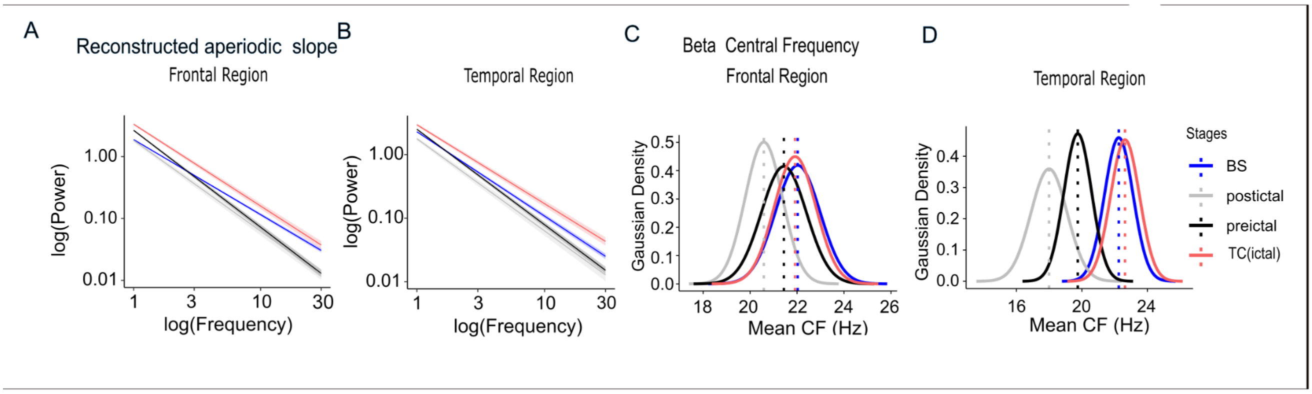


**Fig. S7.** A. Reconstructed aperiodic slope in log–log space, showing steepening during preictal and postictal stages and flattening during the ictal stage relative to preictal and postictal phases in Frontal and B. Temporal brain region. C. Periodic Central Frequency (12–30 Hz) peak in the beta band across baseline, preictal, ictal, and postictal stages in Frontal D. and Temporal regions of Epm2a−/− mice.

**Methods:**

### 2.2 Surgical Protocol, Animal Preparation

All surgical procedures were conducted on both wildtype and transgenic C57BL/6 mice (Laforin knockout model) using aseptic techniques. Mice were anesthetized with isoflurane (5% for induction, 1.5% for maintenance in 100% $O_2$) and secured in a stereotaxic frame ( Stoelting Co.). Anesthetic depth was periodically monitored by confirming the absence of a pedal withdrawal reflex. Body temperature was maintained at 37°C throughout the procedure using a feedback-controlled heating pad. For local analgesia, a subcutaneous injection of lidocaine (0.1 mg/kg) was administered at the incision site. A midline incision was made along the scalp to expose the skull. The skull surface was cleaned, and the stereotaxic landmarks of Bregma and Lambda were clearly identified. Five burr holes were drilled through the skull using a 0.8 mm micro-drill bit, taking care not to penetrate the underlying dura mater. A 1 mm long titanium screw was threaded into each burr hole until it made firm contact with the dural surface.

Four active screw EEG electrodes were implanted stereotactically over the frontal and temporal cortices, with ground and reference electrodes placed over the cerebellum. Coordinates relative to Bregma were: right frontal (AP +2.5 mm, ML +1.3 mm), left frontal (AP +2.5 mm, ML −1.3 mm), right temporal (AP −3.0 mm, ML +2.5 mm), left temporal (AP −3.0 mm, ML −2.5 mm), and reference (AP −6.0 mm, ML 0.0 mm). The lead wires from the screws were soldered to a multi-pin connector, which was then securely fixed to the skull. All implanted electrodes were fixed to the skull with dental glass ionomer cement. The sockets were attached to a wire electrode, and the assembly was secured to the skull by silicon glue. This formed a durable head cap for subsequent recordings. Post-surgery, animals were individually housed and monitored for pain and distress.Animals were allowed a recovery period of 3-5 days before the commencement of electrophysiological recordings.For post-surgical pain management, animals received meloxicam (5 mg/kg).

**2.3 EEG Acquisition**

Electrophysiological recordings were carried out in freely moving mice using the Intan RHS stim/recording system (Intan Technologies, LLC). Animals were connected via a lightweight and flexible tether to minimize movement restriction. All recordings were performed inside a Faraday cage to reduce environmental electrical noise and prevent signal loss during transmission.

Neuronal signals were acquired using the Intan RHS stim/recording system (Intan Technologies) in combination with an RHS32 low-noise amplifier headstage. The data were digitized at a sampling rate of 30 kHz, resampled 2kHz. To suppress power-line interference, a 50 Hz notch filter was applied for online real-time visualization. In addition, real-time band-pass and line-noise filtering were employed to improve the ability to assess neuronal activity during recordings and to assist in determining appropriate probe insertion depth. These filters were used exclusively for online monitoring and did not affect the stored raw data. During acquisition, a hardware reference was applied to further minimize common-mode noise and improve signal quality across all channels.

**2.5 Seizure grading and waveform classification:**

Seizure severity in PTZ-induced mice was evaluated using a modified Racine scale in synchrony with EEG/video recordings (Van Erum et al., 2019)..This scoring system quantifies seizure-related behavioral manifestations, with each stage corresponding to a progressive increase in seizure severity. Scores 0–2 were categorized as partial (focal) seizures and classified as low-grade events. Scores 3–4 represented generalized seizures, characterized by bilateral motor involvement and loss of postural stability classified as high-grade events. Higher scores (5–7) denoted severe tonic–clonic seizures, marked by sustained clonus, tonic posturing, and potential loss of righting reflex. Scoring was performed by trained observers blinded to experimental groups to ensure reliability and reproducibility.For tonic-clonic seizure analyses: Preictal epochs were defined as the 30 s segment immediately preceding electrographic Tonic Clonic seizure onset only, excluding periods containing overt ictal activity.Ictal epochs were selected as 30 s segments occurring during sustained electrographic seizure activity accompanied by corresponding behavioral seizure manifestations.Postictal epochs were defined as the first 30 s segment immediately following Tonic Clonic seizure termination, indicating EEG depression.For low-grade (grade-2) and high-grade (grade-4) seizures, epochs were selected from behaviorally scored events with matching electrographic abnormalities as per revised racine scale. Only epochs clearly belonging to the target stage behaviourally were retained. Epochs overlapping transitional periods between states were excluded to avoid contamination between preictal, ictal, and recovery phases.Where seizure duration was shorter than the required epoch length or contained excessive artifact, that event was excluded from epoch-based analyses.
Pathological waveforms occurring during PTZ-induced seizures were classified into four categories based on established electrographic morphology: spike-wave discharges (SWD), characterized by spike components followed by slow-wave components and associated with non-motor seizure behaviors at lower PTZ doses. Polyspike-burst discharges (PSB), consisting of short trains of irregular spikes and polyspikes; high-amplitude polyspikes (HAPS), characterized by high-amplitude polyspikes and spike-wave discharges

accompanying severe tonic-clonic seizure behavior. low-amplitude high-frequency rhythmic waveforms (LAHFRW), corresponding to long-lasting, low-amplitude spiking often followed by high-frequency oscillations, with spiking frequency that varied across the course of the seizure and was consistently followed by postictal EEG depression.Severe seizure behavior was characterized by these high-amplitude polyspike and spike-wave patterns, with behavioral expressions ranging from clonic convulsions to wild jumping, consistent with the seizure-grading framework used here (Van Erum et al., 2019). An EEG discharge pattern of long-lasting, high-amplitude spiking with poly-spikes followed by high-frequency oscillations was found to coincide specifically with the most severe behavioral convulsions, distinguishing it from shorter, lower-amplitude discharge types associated with milder behavioral categories (Lüttjohann et al., 2009).

#### 2.6 EEG Data Processing and Analysis

All offline data processing and analysis were initiated in Brainstorm software (Tadel et al., 2011), running in MATLAB (version R2024a; The MathWorks Inc., Natick, MA, USA). The continuous raw data from the four cortical electrodes were imported into Brainstorm and re-referenced by subtracting the signal from the cerebellar reference electrode. The data were then filtered using a 1–100 Hz band-pass filter and a 50 Hz notch filter to remove line noise. Following preprocessing, the entire recording was visually inspected by an expert reviewer to identify and mark electrographic seizures.

For quantitative analysis, the continuous data from both baseline and post-PTZ events were segmented into non-overlapping 30-second epochs. Epochs containing significant artifacts or electrographic seizure activity were excluded to ensure that subsequent spectral analysis reflected underlying, non-epileptic brain states.

##### 2.6.1 Fitting Oscillations & One-Over-F' (FOOOF) algorithm

To decompose the power spectral density (PSD) of clean epochs into periodic and aperiodic components, we used the 'Fitting Oscillations & One-Over-F' (FOOOF) algorithm (Donoghue et al., 2020), implemented in Python. This separation is crucial as traditional band-power measures can conflate these two signals (Donoghue et al., 2020). The FOOOF algorithm models the aperiodic component as a 1/f function and identifies periodic components as Gaussian peaks rising above the aperiodic slope.

Power spectral densities were estimated using Welch's method with a Hamming window, as implemented in the Brainstorm toolbox. The FOOOF algorithm was applied to the PSD of each epoch, modeling the spectrum over a frequency range of 1–30 Hz. The restricted frequency range was chosen because higher frequencies showed substantially lower signal-to-noise ratios in screw EEG recordings and because robust parameter estimation was prioritized over direct replication of higher-frequency E/I-sensitive ranges.
FOOOF was applied independently to the power spectral density (PSD) estimated from each non-overlapping 30 s epoch and for each recording channel separately; spectra were not averaged prior to fitting.The algorithm parameters were set as follows: peak width limits = [1.0, 8.0] Hz; maximum number of peaks = 6; minimum peak height = 0.1; peak threshold = 2.0 standard deviations; and aperiodic mode = 'fixed'. From the model output for each epoch, we extracted the aperiodic parameters (offset and exponent) and the periodic parameters (center frequency, power, and bandwidth of each identified oscillatory peak). These extracted parameters were then used for all subsequent statistical comparisons and classification analyses. We selected fixed-mode parameterization because visual inspection

of group-average spectra did not reveal consistent knee-like transitions within the 1–30 Hz fitting range and because the restricted frequency range limits stable estimation of the knee parameter.

The aperiodic exponent has been proposed as a systems-level marker related to excitation and inhibition balance, with computational models and empirical studies suggesting that flatter slopes are often associated with relatively greater excitation and steeper slopes with relatively greater inhibition (Gao et al., 2017; Donoghue et al., 2020; Chini et al., 2022). Both the aperiodic parameters (exponent and offset) and the band-specific oscillatory powers were then used for all subsequent statistical comparisons. All model parameters reported in the manuscript were applied uniformly across all animals, genotypes, channels, and experimental conditions. During preliminary analyses, we evaluated multiple frequency ranges (1–100 Hz, 1–60 Hz, and 1–30 Hz) as well as different epoch durations (2 s, 5 s, 10 s, 20 s, and 30 s) for FOOOF-based parameterization. Based on comparative model performance, 30 s epochs yielded the most reliable goodness-of-fit estimates and were therefore selected for the final analyses, consistent with prior recommendations in the literature. 1-30 Hz frequency was suitable due to screw-EEG SNR and to avoid line-noise contamination.

Interictal Epileptiform Discharge (IED) Detection Raw EEG recordings were processed at their native sampling rates. To eliminate mains powerline interference, a time-domain second-order biquad notch filter was applied at 50 Hz. A per-channel artifact mask was then computed to exclude non-physiological segments from baseline estimation and event detection. Samples were flagged as artifacts if their amplitude exceeded 12 standard deviations from the channel mean or if they belonged to a run of five or more consecutive identical samples (indicating amplifier saturation or clipping). The resulting mask was dilated by 10 ms on each side, and these flagged segments were excluded from downstream analysis.

#### 2.6.3 Time-Resolved Spectral Parameterization (SPRiNT)

Spectral parameterization of the EEG signal was performed using SPRiNT (Spectral Parameterization Resolved in Time; Wilson et al., 2022)., implemented within Brainstorm as process_sprint. SPRiNT extends the FOOOF/specparam framework (Donoghue et al., 2020) to resolve aperiodic and periodic spectral features across time by applying the specparam algorithm to a series of short-time Fourier transform (STFT) windows within each epoch.

For each 30-second epoch, a sliding STFT was computed using windows of 1 second duration with 90% overlap, producing a time-frequency representation with a frequency resolution of 1 Hz. The specparam algorithm was then applied to the power spectrum at each time point within the frequency range of 1–30 Hz. Local temporal averaging was performed across 3 consecutive time-frequency estimates (loc_average = 3) prior to model fitting, in order to improve signal-to-noise ratio of the spectral estimates. Outlier time points were identified based on a maximum frequency deviation threshold of 2.5 Hz and a minimum temporal neighborhood requirement of 3 neighboring estimates within a 4 time-step window were subsequently removed.

The aperiodic component of each power spectrum was modeled using a fixed-exponent parameterization according to the following equation:

$$\log P(f) = b - \log(f ^ \chi)$$

where P(f) is the power spectral density at frequency f, b is the aperiodic offset (intercept), and χ is the aperiodic exponent (slope). The offset reflects the overall broadband power level of the signal, while the exponent characterizes the steepness of the 1/f-like spectral decay and has been linked to the excitation–inhibition (E:I) balance of the underlying neural circuitry (Gao et al., 2017; Donoghue et al., 2020).

Following removal of the aperiodic component, periodic peaks were identified in the flattened spectrum as Gaussian functions. Peak detection was constrained by the following parameters: center frequency within 1–30 Hz, peak width between 0.5 and 12 Hz, a minimum peak height of 3 (in log power units above the aperiodic fit), and a maximum of 3 peaks per time point. A proximal threshold of 2 Hz was applied to prevent detection of overlapping peaks. Model fitting was performed using least-squares optimization. The goodness-of-fit of the full parameterization model at each time point was assessed using the mean squared error (MSE) between the modeled and empirical log power spectra.

#### 2.6.5 Data Visualization

For aperiodic parameters (exponent and offset), data were visualized using box plots with notches representing the 95% confidence interval around the median. A yellow diamond-shaped marker indicated the mean value for each condition.

For band-specific oscillatory parameters, center frequencies (CF) were visualized using Gaussian density curves. For each experimental stage, the mean center frequency and mean bandwidth (STD) across all detected peaks within that band were calculated. These summary statistics were used to generate normal distribution curves with the mean CF as the peak location and the mean bandwidth as the spread. Vertical dotted lines marked the mean CF for each stage. This visualization approach represented the average spectral characteristics of oscillatory activity within each frequency band while accounting for inter-epoch variability.

Oscillatory power (amplitude extracted from FOOOF) were visualized using box plots with the same format as the aperiodic parameters notches representing the 95% confidence interval around the median and a yellow diamond-shaped marker indicating the mean value for each condition.

### 2.7 Statistical Analysis

All statistical analyses were conducted using Bayesian multilevel regression models implemented in the brms package (Bürkner, 2017) in R (version 4.2.1; R Core Team, 2022). Bayesian approaches were chosen for their ability to account for hierarchical data structures, provide full posterior distributions for parameter estimates, and quantify uncertainty in a probabilistic framework (Kruschke, 2015). We used separate comparison group-wise since not all animals had grade-4 or TC after the same dose of PTZ. So changes during different grades were compared to baselines of the same subject to rule out group level effects.

All models included random intercepts for animal_ID to account for inter-animal variability. Models were fit using 4 chains with 2000 iterations each (1000 warmup), and convergence was assessed via R-hat values (all < 1.01). A fixed random seed (123) was used to ensure reproducibility. The specific model structures used for different analyses (see Model List). In general, models included experimental conditions as fixed effects with appropriate interaction terms, following the general form:

Outcome ~ Predictor(s) + (1 | animal_ID)

##### 2.7.1 Effect Size Calculation

For all models, effect sizes were calculated as standardized coefficients by dividing each Stage coefficient (β_Stage) by the residual standard deviation (σ) from the model:Cohen's d = β_Stage / σ. This approach yields standardized effect sizes representing the magnitude of stage differences relative to within-unit residual variation, independent of measurement scale. These standardized coefficients were summarized using posterior means and 95% credible intervals.

##### 2.7.2 Model Validation

Posterior predictive checks were conducted for all models using the pp_check() function from BRMS to assess model fit by comparing observed data to data simulated from the posterior predictive distribution. This ensured that the models adequately captured the data-generating process.

Model List

Baseline (BS) - Isoflurane (IF) - Wild Type (WT)

- Exponent: exponent ~ Stage × region_group + (1 | animal_ID)
- Offset: offset ~ Stage × region_group + (1 | animal_ID)
- Center Frequency (CF): CF ~ Stage + (1 | animal_ID)

Baseline (BS) - Isoflurane (IF) - Laforin Knockout (LKO)

- Exponent: exponent ~ Stage + (1 | animal_ID)
- Offset: offset ~ Stage + (1 | animal_ID)

Baseline (BS) - Awake Recordings Only

- Exponent: exponent ~ group + (1 | animal_ID)
- Offset: offset ~ group + (1 | animal_ID)
- Amplitude: Amp ~ group × region_group + (1 | animal_ID)

Tonic-Clonic (TC) Seizure Stages

- Exponent: exponent ~ Stage × region_group + (1 | animal_ID)
- Offset: offset ~ Stage × region_group + (1 | animal_ID)

Notes:

- × denotes interaction terms between predictors
- Stage refers to experimental stages (baseline, preictal, tonic-clonic, postictal, grade-2, grade-4)
- group refers to genotype (LKO vs WT)
- region_group distinguishes recording sites (Left/ Right, Frontal, Temporal)
- All models include random intercepts for animal_ID

**2.7.3 Hurst Exponent Analysis:** Hurst Exponent Analysis was done to quantify the fractal nature and long-range temporal dependency (memory) of the electrophysiological signals, the Hurst exponent (H) was computed channel-wise across each recording. Electrophysiological data were analyzed at their native sampling rate of 2 kHz and segmented into consecutive, non-overlapping 2-second windows. The use of non-overlapping windows was selected to guarantee statistical independence between samples, following established EEG and local field potential (LFP) analysis protocols (Geng et al., 2011; Witton et al., 2019). For each window and channel, H was estimated via Rescaled Range (R/S) analysis using the PyEEG package. The resulting exponent describes signal persistence: H approx 0.5 indicates a random walk (white-noise-like, memoryless), $H > 0.5$ indicates persistent (trending, self-similar) dynamics, and $H < 0.5$ indicates anti-persistent (mean-reverting) dynamics.

List of Abbreviations used:
E/I: Excitation/Inhibition
EEG: Electroencephalogram
FOOOF: Fitting Oscillations & One-Over-F
LKO: Laforin Knockout
PSD: Power Spectral Density
WT: Wild-Type
GABA: Gamma-aminobutyric acid
AED: Antiepileptic Drug
M/EEG: Magneto/Electroencephalography
MEG: Magnetoencephalography
IAEC: Institutional Animal Ethics Committee
$O_2$: Oxygen
AP: Anteroposterior
ML: Mediolateral
RHS: RHD2000 series recording system
PTZ: Pentylenetetrazol
i.p.: Intraperitoneally
LDA: Linear Discriminant Analysis
TC: Tonic-Clonic
TLE:Temporal Lobe Epilepsy
CI: Confidence Interval
LF: Left Frontal
RF: Right Frontal
LT: Left Temporal
RT: Right Temporal
IF: Isoflurane
CF: Central Frequency
IEDs: Interictal Epileptic Discharges
SWDs: Spike-and-Wave Discharges
ECoG: Electrocorticography
REMS: Rapid Eye Movement Sleep
